# Diagnosability to inform species delimitation for the genus Emydura (Testudines: Chelidae) from northern Australia

**DOI:** 10.1101/2025.07.10.664252

**Authors:** Arthur Georges, Peter J. Unmack, Andrzej Kilian, Xiuwen Zhang, Yolarnie Amepou, Duminda S.B. Dissanayake

## Abstract

Understanding the evolutionary history of diversifying lineages and the delineation of species remain major challenges for evolutionary biology. Here we use single nucleotide polymorphisms (SNPs) and sequence fragment presence-absence (SilicoDArT) data to combine phylogenetics and population genetics to assess species boundaries with a focus on diagnosability. We challenge current and proposed taxonomies in a genus of Australian freshwater turtles (Chelidae: *Emydura*) from northern Australia and southern New Guinea. In a six-step process, we combine phylogeny with the concept of diagnosability based on fixed allelic differences to select diagnosable lineages as candidate species. Four taxa are supported as diagnosable lineages, two of which we elevate to species status. The nuclear and mitochondrial phylogenies differed in important respects, which we attribute to recent or contemporary lateral transfer of mitochondria during hybridization events, deeper historical hybridization or possibly incomplete lineage sorting of the mitochondrial genome. Taxonomic decisions in cases of allopatry require subjective judgement. Our six-step strategy and the necessary (but not sufficient) criterion of diagnosability adds an additional level of objectivity before that subjectivity is applied, and so reduces the risk of taxonomic inflation that can accompany lineage approaches to species delimitation.

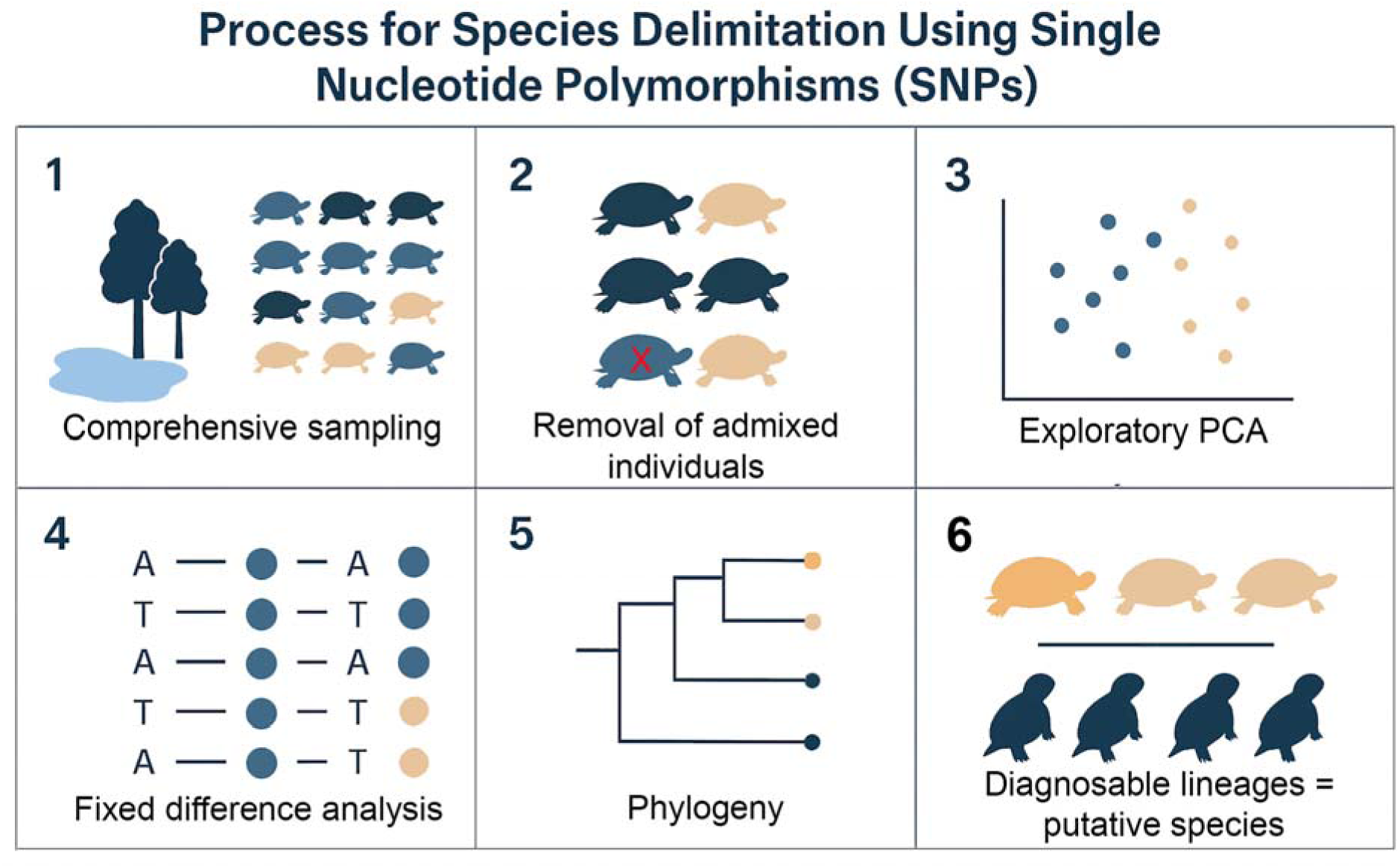

## INTRODUCTION

Speciation, in its simplest conception (Dobzhansky, 1951, Mayr, 1963), involves the progressive divergence of two lineages in the absence of geneflow until reproductive compatibility is eroded and ultimately the two lineages become irrevocably committed to distinct evolutionary fates. Early in this process, isolation of two sister lineages may not be complete, in which case speciation begins with a pattern of reduced introgression in the regions of the genome that are most differentiated between two divergent lineages (Gompert *et al*. 2012, Harrison and Larson 2016). It ends when reproductive compatibility becomes compromised (Coyne and Orr 1989, Orr 1995), giving permanence to the genetic isolation formerly arising from potentially ephemeral geographic isolation.

This process of allopatric speciation is accompanied by accumulation of measurable differences in allelic profiles of progressively diverging lineages and gene pools. Early in the process, differences in frequencies of alleles, though still held in common, will accrue among populations. This will be followed by some alleles coming to fixation in one population and the accumulation of private alternate alleles in others, under drift (Wright 1931, Jensen *et al*. 2019) or selection (McDonald and Kreitman 1991). Ultimately, fixed allelic differences between populations will arise, where two populations share no alleles at a locus, though not necessarily uniformly across the genome (Ellgren *et al*. 2012, Cruickshank and Hahn 2014). In the biallelic state, such as is characteristic of single nucleotide polymorphic markers (SNPs), a fixed difference occurs when one allele at a locus becomes fixed in one population and the alternate allele becomes fixed in the other population. Unlike allele frequency differences, fixed allelic differences are irreversible in the absence of parallel mutations and in the absence of exchange of lost alleles through low or episodic geneflow. Parallel mutations, that is, multiple mutations at a single nucleotide site, are thought to be rare in SNP data at shallow levels of divergence (Brumfield *et al*. 2003) and unlikely in any case to move to fixation unless under intense selection (Sterns 2013). As such, acquisition of a fixed allelic difference is a significant biological event (Richardson *et al*. 1986). Fixed allelic differences are often viewed as a key indicator that populations have diverged into separate species or evolutionarily significant units (Templeton, 1989, Coyne and Orr 2004, Wiens and Penkrot 2002).

Phylogenies built from genetic profiles at the individual or population level capture the accumulation of genetic differences in the form of divergent lineages and the nested bifurcating structure that we take to represent the patterns of ancestry and descent among deeper lineages and their descendent clades. The substantive lineages to emerge from a phylogenetic analysis can be either species or represent population structure within species. Biodiversity assessment and monitoring for conservation does not require a decision on which lineages are species and which are lineages within species – diversity can be measured as phylogenetic diversity with an appropriate weighting for level of divergence (Faith 1992, Manson *et al*. 2022). Nevertheless, the delineation of species remains important (Sites and Marshall 2003). Species are not simply constructs for classification as might be argued for higher level taxonomic levels, but are real biological entities (Hillis *et al*. 2021). The genetic and demographic processes by which species arise (speciation) or are extinguished (extinction) remain subjects of rigorous enquiry. The formal description of species remains important for governments to set conservation priorities and fund conservation initiatives (Pante *et al*. 2015). For these reasons, species delimitation also remains a subject of intense interest.

Making a decision on which lineages in a phylogeny should be regarded as species and which represent lineage diversity within species requires judgement in all cases but those involving sympatric taxa. Such judgements clearly depend upon which species concept is adopted. The biological species concept which requires indirect inferences on reproductive isolation in allopatry (Mayr, 1963) and a lineage species concept which has every substantive lineage as a species (e.g. Fujita *et al*. 2012), are perhaps at two ends of the decision spectrum. In this paper, we take an intermediate view by arguing that diagnosability is a necessary (but not sufficient) operational criterion for regarding a lineage as a species. A diagnosis is the foundation of species descriptions using morphology (Rheindt *et al*. 2023) dating back to the time of Linneaus (Renner 2016). A diagnosis is constructed from a set of characters that, alone or in combination, are able to distinguish the focal taxon from other taxa. In molecular studies, the focus on the taxon in a definition of diagnosability, such as that adopted by the ICZN (Rheindt *et al*. 2023), admits the possibility that two taxa can be distinguished on the basis of their allele frequency profiles, represented for example in a Principal Components Analysis (PCA, Jolliffe 2002). This leaves ambiguous the assignment of some or even many individuals to one taxon or another. We take the view in this paper that diagnosability implies that one should be able to, on the basis of diagnostic markers or traits, reliably assign every individual to its species. An evolutionary lineage – a linear series of ancestral and descendant populations or metapopulations and their descendant clade – is a putative species if all the individuals assigned to a lineage can be distinguished from all individuals of other lineages by one or more diagnostic characters.

There is an attendant benefit of applying the criterion of diagnosability in population genetics. Lack of diagnosability manifests as shared alleles at all loci. Lack of diagnosability means one cannot confidently reject the null proposition that two putative lineages are on an interconnected evolutionary trajectory – that the two putative lineages are (or have recently been) subject to geneflow and admixture. Thus, diagnosability applies a filter to lineages, retaining only those for which there is evidence of isolation in the accumulation of fixed differences, either by reproductive or geographic isolation. Such lineages are candidates to be considered either as species or as diagnostic lineages within species. Other branches within the phylogeny, however well supported by bootstrap values, need not be considered further in the context of species delimitation.

High throughput parallel sequencing (next generation sequencing, Metzker 2010) and low cost reduced representational sampling of the genome for single nucleotide polymorphisms (SNPs) – DArTSeq, RADSeq, ddRAD (Jaccoud *et al*. 2001, Baird *et al*. 2008, van Tassell *et al*. 2008, Sansaloni *et al*. 2011, Kilian *et al*. 2012, Peterson *et al*. 2012) – have enabled genomics at the level of populations to be considered with phylogenomics in studies of the pattern and process of speciation, species delimitation and phylogeography (Georges *et al*. 2018). Genotyping of geographically comprehensive samples each with adequate replication is now possible to deliver better understanding of the historical and contemporary drivers of geographic patterns in genetic diversity at regional scales involving species, subspecies, substantive lineages and other evolutionarily significant units.

Our empirical approach has been presented in earlier papers (Adams *et al*. 2014, Georges *et al*. 2018, Unmack *et al*. 2022). It is applied here with variations. There are six steps to our analysis. First, we require comprehensive coverage of populations within the target taxa, both to avoid interpreting sparse sampling of demes on a cline as distinct taxa (Marshall *et al*. 2021) and to capture evidence of any geneflow across contact zones. It also requires sampling of sufficient intensity at each collection site as to adequately characterize local allelic profiles (Adams *et al*. 2022). Second, we require that instances of recent and contemporary hybridization and introgression between putative taxa be identified. Establishing diagnosability, whether it be based on morphological data or genetic data, can not be achieved if we include hybrids between the two putative taxa being compared. In the case of SNP data, a single F1 hybrid, easily detected, would be sufficient to obliterate any otherwise fixed allelic differences between two putative taxa. We identify and set aside hybrids and backcrosses qualitatively by examination of ordination plots and quantitatively using the software NewHybrids (Anderson and Thompson 2002). The third step is exploratory, using PCA to identify structure among the sampled individuals and, in particular, to identify any putative groupings based on differing allelic profiles. This exploratory step sets the expectation for the analyses to follow, allows the subsequent identification of clusters of individuals that differ in allelic profiles without being diagnostic, and provides a visual representation of potential clines that may later come to be progressively aggregated in a fixed allelic difference analysis. The fourth step is to identify diagnosable aggregations (sensu Templeton, 1989) using analysis of fixed allelic differences, recursively amalgamating populations that lack fixed differences to yield a set of diagnosable aggregations (Georges *et al*. 2018). Fifth, we apply phylogenetic techniques to the sampled populations (typically sample sites) to identify well supported lineages. There is a diversity of approaches that can be taken here. The primary focus of this paper lies in the sixth and final step, where we identify the lineages in the phylogeny that are diagnosable by bringing together the diagnosable aggregations of individuals arising from the fixed difference analysis and the lineage structure identified in the phylogenetic analysis. Those lineages that represent diagnosable aggregations are either candidate species or represent substantive diagnosable lineage structure within species (Evolutionarily Significant Units, ESUs). A decision between the two requires subjective judgement where possible, drawing upon other available data, morphological, physiological, behavioural.

We apply this empirical approach to the delimitation of species of freshwater turtle (genus *Emydura* Bonaparte, 1836) widely distributed across northern Australia and southern New Guinea. This complements an earlier study of the southern *Emydura* (Georges *et al*. 2018). The range of the northern species of *Emydura* extends the full extent of the north of the Australian continent (2,300 km) and the south of New Guinea (1,500 km) from west to east. They are obligate freshwater organisms with dispersal constrained by the ocean, well-defined drainage divides and dendritic riverine structure. Their taxonomy has a complex history (reviewed by Cann and Sadlier 2017, Kehlmaier *et al*. 2024 and I. Smales, in preparation). Today, three species of *Emydura*, one represented by two distinct subspecies, are accepted as occurring in northern Australia and southern New Guinea, though competing taxonomies exist (Cann and Sadlier 2017, TTWG 2017). These three northern species include: the northern redfaced turtle *Emydura australis* (Gray, 1841) (Figure 1a,b) whose range extends across the rivers draining the Kimberley Plateau in north Western Australia to the Daly River of the Northern Territory; the diamondhead turtle *Emydura subglobosa worrelli* (Wells and Wellington, 1985) (Figure 1c,d) from the rivers draining into the Gulf of Carpentaria and Arnhem Land; the New Guinea painted turtle *Emydura subglobosa subglobosa* (Krefft, 1876) (Figure 1e,f) widely distributed across New Guinea south of the central ranges and in the Jardine River at the tip of Cape York in Australia; and the northern yellowfaced turtle *Emydura tanybaraga* Cann, 1997 (Figure 1g-i) with a poorly defined but disjunct distribution in the Northern Territory and northern Queensland. Populations within these species, in Australia at least, exhibit great variability across their ranges in colour pattern and shell shape (Cann and Sadlier 2017). The reference to species as redfaced and yellowfaced has led to considerable confusion because the character is unreliable (refer Cann and Sadlier 2017:379, second figure) and this trait does not correlate well with mitochondrial genotype (Kehlmaier *et al*. 2024). The identification of *Emydura tanybaraga* in the field has been particularly problematic, confusion that has been made worse by hybridization between the taxa where their distributions overlap and resultant mixed signals in their morphological identification.

**Figure 1.**
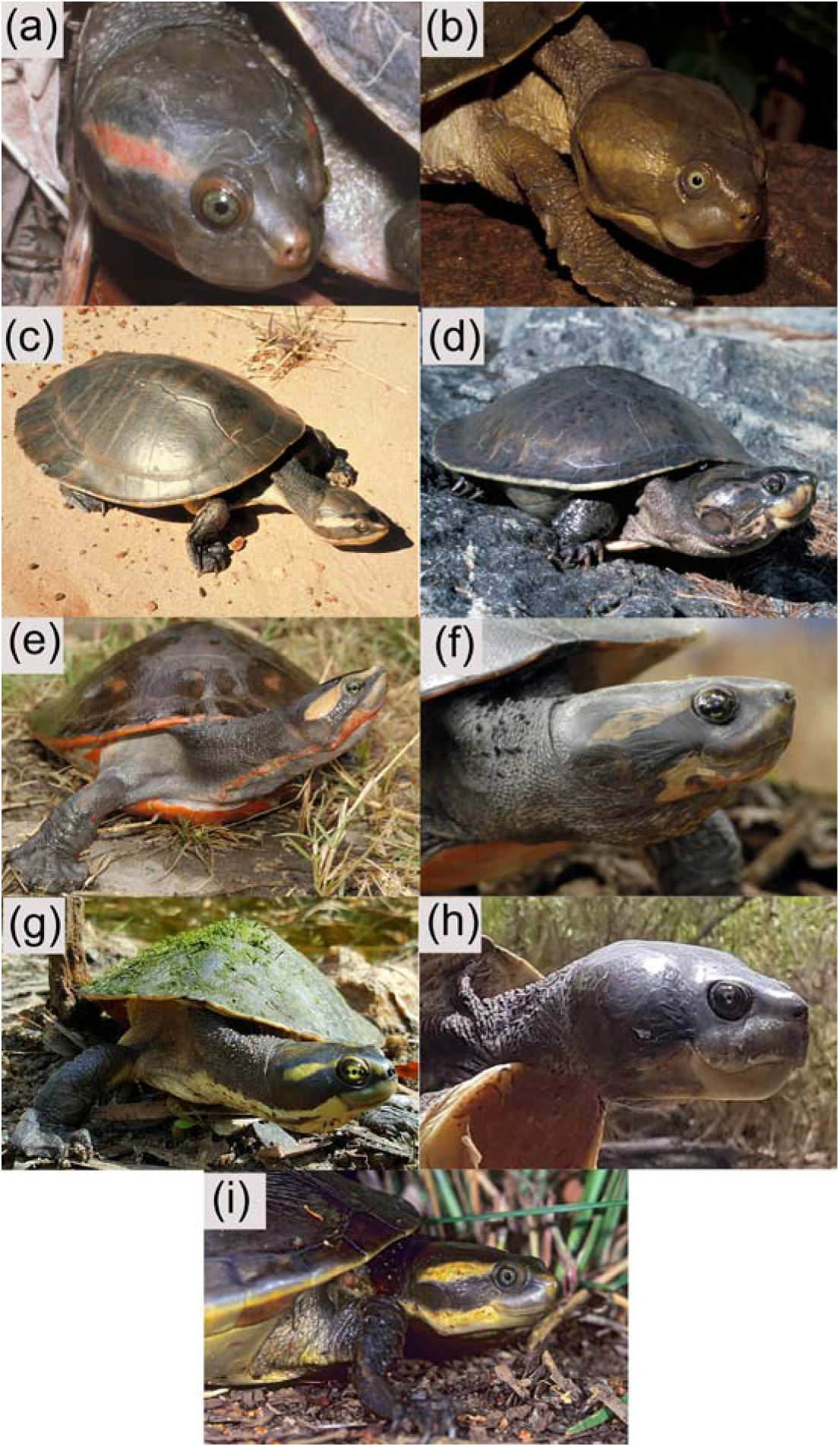
Species and subspecies of *Emydura* from northern Australia and Southern New Guinea. (a) *Emydura australis*, Jasper Gorge, Victoria River NT; (b) *E. australis*, aged individual, Daly River NT; (c) *E. subglobosa worrelli*, NT [Photo: Conservation Commission of the NT); (d) *E. s. worrelli*, aged specimen, Roper River NT; (e) *E. s. subglobosa*, Suki-Aramba swamps, Fly River PNG; (f) *E. s. subglobosa*, aged specimen, Suki-Aramba swamps, Fly River PNG; (g) *E. tanybaraga*, Archer River, Qld [Photo: Jason Schaffer]; (i) *E. tanybaraga*, aged individual, Archer River, Qld [Photo: Jason Schaffer]; (i) specimen of a form provisionally assigned to *E. tanybaraga*, Mitchell River WA [Photo: Jiri Lochman, -14.8366, 125.6291, 153 m ASL]

Here, we use SNP, SilicoDArT and mitochondrial sequence data to evaluate species boundaries of the northern *Emydura*. We interpret our results in the context of the controversy on the use of lineages (and their resultant clades) to delineate species (Hoelzel 2016, Sukumaran and Knowles 2017) and inject the concept of diagnosability of lineages as a necessary but not sufficient criterion for selecting lineages as putative species. Our study provides strong support for the recognition of the existing taxa rather than new or alternative classifications. We elevate *Emydura subglobosa subglobosa* and *Emydura subglobosa worrelli* to full species. There was considerable discordance between the phylogenies based on nuclear SNP/SilicoDArT data and the mitochondrial phylogeny, which we attribute to lateral transfer of mitochondria between *Emydura tanybaraga* and the other taxa.

## 2 MATERIALS AND METHODS

### 2.1 Specimen Collection

For the nuclear marker genotyping, 526 individuals were sampled from 43 drainage basins of northern Australia from the Fitzroy River in the northwest (Western Australia) to the Pascoe River of Cape York in the northeast (Queensland) and Papua New Guinea south of the central ranges (Figure 2). This sampling covers the full range of the species currently regarded as *Emydura australis*, *Emydura subglobosa* (subspecies *subglobosa* and *worrelli*) and *Emydura tanybaraga* but excludes detailed spatial coverage of the southern *Emydura macquarii* (Gray, 1830) (subspecies *emmottii*, *gunnabarra*, *krefftii*, *nigra*, and *macquarii*), the latter having been dealt with in an earlier paper (Georges *et al*. 2018). Our sampling target was 10 individuals per population (not always achieved). Trees generated by the nuclear SNP and SilicoDArT analyses are unrooted (but see Georges and Adams, 1992, 54 allozyme loci) and for the mitochondrial analysis were rooted with *Elseya flaviventralis* Thomson and Georges 2016 (after Kehlmaier *et al*. 2024 based on both nuclear and mitochondrial data). A full list of specimens and localities is provided in supplemental information (Tables S1-S3). Our taxonomy follows that of Georges and Thomson (2010) with the change of *Emydura victoriae* (Gray, 1842) to *Emydura australis* and clarification of the holotypes for both by Kehlmaier *et al*. (2024). Nomenclature for drainage basins follows that of the Australian Drainage Divisions and River Basins (Auslig 2001) with the separation of the Carson and Mitchell rivers from the King Edward River drainage, Western Australia (WA), and other minor changes (e.g. we use McKinlay River which is in the Mary River drainage of the NT). Where there is ambiguity, we add the state abbreviation to the river name, e.g. Mitchell River WA (Western Australia) or Mitchell River Q (Queensland).

**Figure 2.**
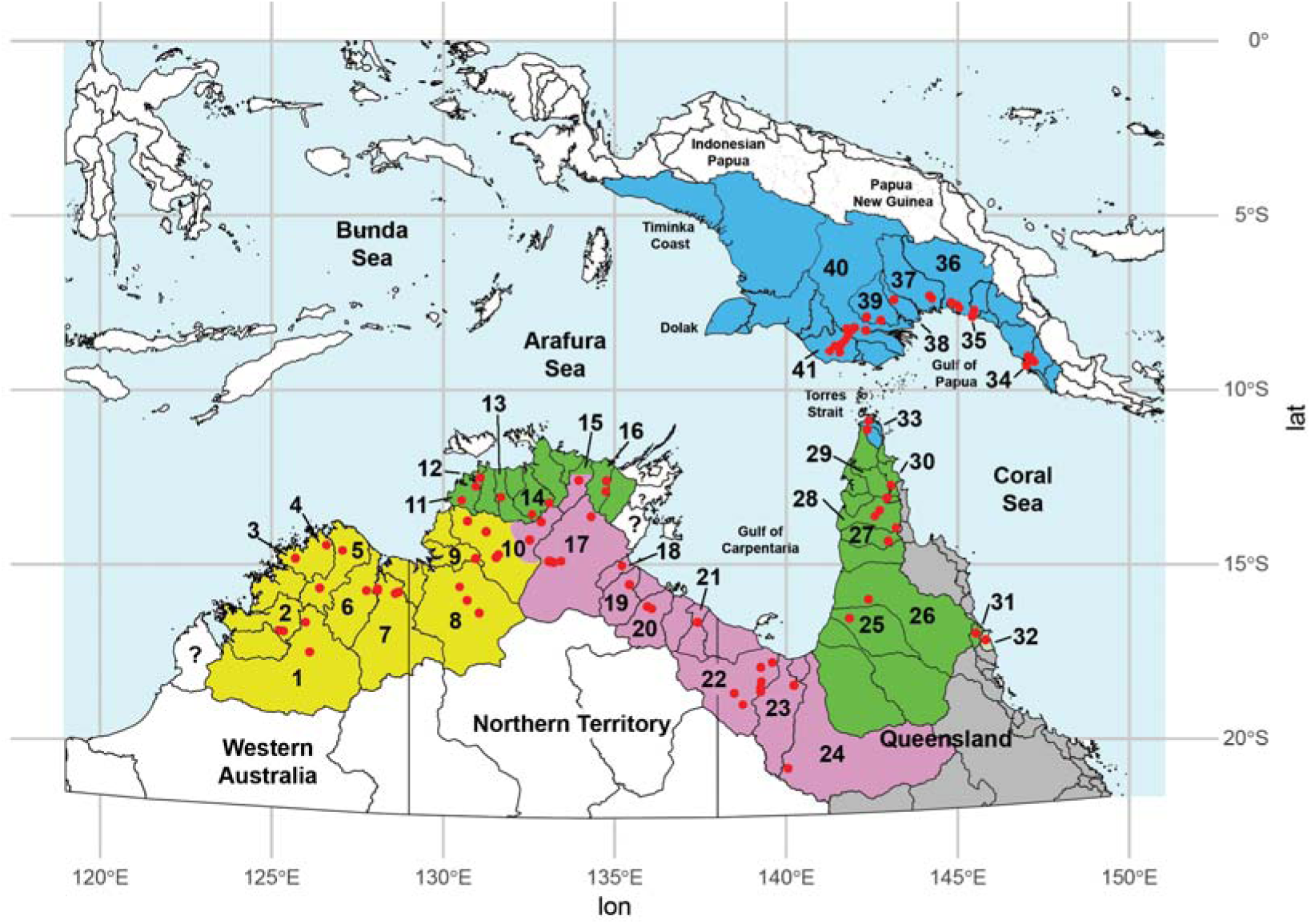
Drainage basins from which samples were taken for the SNP and SilicoDArT analyses. *Emydura australis* (yellow): 1, Fitzroy R WA (n=11); 2, Isdell R (n=12); 3, Mitchell R WA (n=4); 4, Carson R (n=10); 5, Drysdale R (n=10); 6, Pentecost R (n=5); 7, Dunham R (n=5) & Ord R (n=15); 8, Victoria R (n=16), 9, Fitzmaurice R (n=8); 10, Daly R (n=64). *Emydura tanybaraga* (green): 10, Daly R (n=10); 11, Finniss & Reynolds rivers (n=2); 12, Darwin (n=1) & Howard rivers (n=5); 13, McKinlay R (n=2); 14, South Alligator R (n=1); 16, Blyth R (n=12); 25, Staaten R (n=6); 26, Mitchell R Qld (n=13); 27, Holroyd R (n=10); 28, Archer R (n=4); 29, Wenlock R (n=10); 30, Olive-Pascoe R (n=9); 31, Barron R (n=2); 32, Mulgrave-Russell R (n=2). Specimens of *E. tanybaraga* occurred with *E. australis* in the lower reaches of the Daly River, but are not shaded. *Emydura subglobosa worrelli* (pink): 10, Daly R (n=25); 15, Liverpool R (n=10); 17, Roper R (n=21); 18, Towns R (n=1); 19, Limmen-Bight R (n=10); 20, McArthur R (n=8); 21, Calvert R (n=10); 22, Nicholson R (n=34); 23, Leichhardt R (n=10); 24, Flinders R (n=5). *Emydura subglobosa subglobosa* (blue): 33, Jardine R (n=16); 34, Vanapa R (n=21); 35, Vailala R (n=10); 36, Purari R (n=3); 37, Kikori R (n=5); 38, Tourama R (n=1); 39, Bamu-Aramia R (n=26); 40, Fly River (n=55); 41, Morehead (n=7) & Bensbach rivers (n=4). Note that drainages 1-7 drain the Kimberley Plateau; drainages 10, 13-17 drain the Arhnem Land Plateau; the Olive-Pascoe [30], Barron [32] and Mulgrave-Russell rivers drain to the east of the Great Dividing Range. Grey shaded drainages are occupied by the southern *Emydura*, *E. macquarii*. The ranges shown here are new, informed by the results of the current work.

Tissue samples comprised a small sliver of skin tissue taken from the trailing edge of the clawless toe of the hind foot in most cases, preserved in 95% ethanol and stored at -20°C, or frozen in liquid nitrogen and stored at -80°C. In some cases, blood was sampled from the jugular vein using a 23-gauge needle and syringe, also preserved in 95% ethanol and held at -20°C. A few samples, drawn from earlier studies (Georges and Adams, 1992, 1996), comprised muscle or blood, snap frozen in liquid nitrogen and stored at -80°C.

### 2.2 DNA Extraction and Sequencing

DNA was extracted by Diversity Arrays Technologies (DArT Pty Ltd, Canberra, Australia) using a NucleoMag 96 Tissue Kit (Macherey-Nagel, Düren, Germany) coupled with NucleoMag SEP (Ref. 744900) to allow automated separation of high quality DNA on a Freedom Evo robotic liquid handler (TECAN, Männedorf, Switzerland). Tissue was first incubated for four hours (blood) or overnight (skin) with proteinase K, adjusted in concentration depending on the tissue.

Sequencing for SNP genotyping was done using DArTseq™ (DArT Pty Ltd) which uses a combination of complexity reduction using restriction enzymes, implicit fragment size selection and next generation sequencing (Sansaloni *et al*. 2011), as described in detail by Kilian *et al*. (2012). The technique is similar to double-digest restriction associated DNA sequencing (ddRAD) (Peterson *et al*. 2012) but has the advantages of accepting lower quantities of DNA, greater tolerance of lower quality DNA and yielding lower allelic dropout rates (Sansaloni *et al*. 2011). The restriction enzyme combination of PstI (recognition sequence 5′-CTGCA|G-3′) and SphI (5’-GCATG|C-3’) was selected on the basis of the evaluation undertaken by Georges *et al*. (2018).

DNA samples were processed in digestion/ligation reactions but replacing a single PstI-compatible adaptor of Kilian *et al*. (2012) with two different adaptors annealed to the two restriction enzyme overhangs. The PstI-compatible adapter included the Illumina flow cell attachment sequence, a sequencing primer sequence, a barcode region of variable length (see Elshire *et al*. 2011) and the PstI-compatible overhang sequence. The reverse adapter contained flow cell attachment sequence and SphI-compatible overhang sequence. Only fragments generated by the PstI-SphI double digest were effectively amplified in 30 rounds of polymerase chain reaction (PCR). Amplifications consisted of an initial denaturation step of 94°C for 1 min, followed by 30 cycles of PCR with the following temperature profile: denaturation at 94°C for 20 s, annealing at 58°C for 30 s, and extension at 72°C for 45 s, with an additional final extension at 72°C for 7 min. After PCR, equimolar amounts of amplification products from each sample were pooled and applied to c-Bot (Illumina) bridge PCR for sequencing on the Illumina Hiseq2500. The sequencing (single end) was run for 77 cycles.

Genomic DNA was extracted for whole mitochondrial sequencing using a salting out method (FitzSimmons *et al*. 1995) or a Qiagen DNeasy blood and tissue kit (Qiagen Australia, Doncaster, Vic, Australia). Three overlapping fragments (ranging in length from 5 to 8 kb) for each mitogenome were amplified using primer pairs CEE12sF (5’-TACAAACTGGGATTAGATACCCCACTATGC-3’) – CEE8354R (5’ ACCCCTAATGATGGTACTGCTCATGAGTGT-3’; CEE8003F (5’ ACCAGTACAATAGATGCCCAAGAAGTAGAAA-3’) – CEECytBR (5’ TTAGCAGGTGTAAAGTTGTCTGGGTCTCCT-3’); CEEND4F3 (5’-CAAACATTAGACTGTGGATCTAAAAATAGGAGTTAAA-3’) – CEE12sR2 (5’-CTCAGTTGGCTACACCTTGACCTGACTT-3’). PCRs used Ranger DNA polymerase (Bio-21117, Bioline/Meridian Australia) under the following conditions: 50–200 ng of genomic DNA, 10 μM of each primer, 5 μL of 5×buffer supplemented and 2.0 U of Ranger DNA polymerase in a final volume of 25 μL. The reactions were performed with an initial denaturation step at 95°C for 2 min; 35 cycles of denaturation at 95°C for 15 s, annealing and extension at 72°C for 8 min and a final extension at 72°C for 10 min. The PCR products were purified with Qiaquick PCR purification kit (Qiagen, Australia) and shipped to BGI (Shenzhen, China) for sequencing. DNA libraries were constructed directly from 1000 ng of purified PCR products following the general library preparation and sequencing protocol to run on an Illumina Hiq2000 sequencer. 100 M reads were obtained for each sample. Sequences were assembled into mitochondrial genomes using *de novo* assembly function in Geneious R6 (Biomatters, Ltd., Auckland, New Zealand) and contigs carrying mitochondrial genomes were identified against all BLASTX database. The whole mitochondrial sequences were complemented with those from Kehlmaier *et al*. (2024) including from relevant holotypes, aligned in Geneious Prime 2024.0.7 (http://www.geneious.com) using Clustal Omega 1.2.2 (Sievers *et al*. 2011).

To expand our coverage of locations across the range of the focal taxa, mitochondrial sequences for Cytochrome B (*cytB*) were generated by DArT Pty Ltd. DArTmp used a 2 step PCR process. Primers were designed to amplify a region of the *cytB* gene while adding a proprietary universal tail sequence, the specific product of the first reaction was used in a second PCR reaction to apply appropriate barcodes to identify samples. The specific primer sequence component used for the *cytB* amplification are: FWD 5’ AACCAYCATTGTYATTCAACTAC 3’, REV 5’ TTAAACTACTGAAATATTGRTT 3’. Amplicons were produced by the first PCR using the following conditions: Kapa PCR Mix 2x (5 µl) (Sigma-Aldrich, Melbourne), specific primer pool 2.5 uM (3 µl), Template (2 µl). The PCR program was set as follows, cycled initially 95°C for 3 min; 29 cycles of 98°C for 20 s, 60°C for 15 s, 72°C for 30 s; 72°C for 2 min; stored at 10C. Amplicons were produced by the second PCR using the following conditions: Buffer MyTaq Ready Mix (2 µl) (Millenium Science, Mulgrave, Australia), 10 µM common primer (2 µl), 2.5 uM barcode primer, MyTaq 5 units/µl (0.02 µl), Template PCR 1 (2 µl), brought to a total volume of 10 µL by addition of water. The second PCR was cycled as follows, 94°C for 30 s, 29 cycles of 94°C for 10 s, 58°C for 30 s, 72°C for 15 s followed by 72°C for 2 min; stored at 10°C. Sequences were generated on an Oxford Nanopore Minion following the protocol for the ligation kit 110 and loaded onto an R9 flowcell.

Resultant *cytB* sequences were complemented with those from Kehlmaier *et al*. (2014), aligned in Geneious Prime 2024.0.7 (http://www.geneious.com) using Clustal Omega 1.2.2 (Sievers *et al*. 2011) and trimmed of the first 72 bp because of multiple gaps and missing bases. Sequences with multiple ambiguous base calls were removed. Coding sequences were checked for unexpected frame shift errors or stop codons using Geneious Prime. Preliminary evaluation of resultant trees was undertaken with a neighbor joining tree and duplicate sequences from the same sample (technical replicates) were identified and removed. Unique haplotypes were identified in DNAcollapser of FaBox 2024.0.7 (http://www.geneious.com, Villisen 2007).

### 2.3 SNP Genotyping

Sequences generated from each lane were processed using proprietary DArT Pty Ltd analytical pipelines. Poor quality sequences were first filtered, applying more stringent selection criteria to the barcode region compared to the rest of the sequence (minimum barcode Phred score 30, pass percentage 75; minimum whole-read Phred score 10, pass percentage 50). In that way, assignment of the sequences to specific samples in the sample disaggregation step was very reliable. Approximately 2,000,000 (±7%) sequences per sample were identified and used in marker calling. These sequences were truncated to 69 bp (including some adaptor sequence where the fragments were shorter than 69 bp) and aggregated into clusters by the DArT fast clustering algorithm, taking advantage of the fixed fragment length, with a Hamming distance threshold of 3 bp. The sequences were error-corrected using an algorithm that corrects a low-quality base (Phred score < 20) to a corresponding high-quality singleton tag (Phred score > 25); where there was more than one distinct high-quality tag, the sequence with the low-quality base was discarded. Identical sequences were then collapsed. These error-corrected sequences were analyzed using DArT software (DArTsoft14) to output candidate SNP markers. In brief, SNP markers were identified within each cluster by examining parameters calculated for each sequence across all samples – primarily average and variance of sequencing depth, the average counts for each SNP allele, and the call rate (proportion of samples for which the marker is scored). Where three sequences survived filtering to this point, the two variants with the highest read depth were selected. The final average read depth per locus was 30.3*x*. One third of samples were processed twice from DNA, using independent adaptors, to allelic calls as technical replicates, and scoring consistency (repeatability) was used as the main selection criterion for high quality/low error rate markers. The DArT analysis pipelines have been tested against hundreds of controlled crosses to verify mendelian behaviour of the resultant SNPs as part of their commercial operations.

The resultant dataset contained the SNP genotypes and various associated metadata of which CloneID (unique identity of the sequence tag for a locus), repAvg (proportion of technical replicate assay pairs for which the marker score is identical), avgPIC (polymorphism information content averaged over the reference and alternate SNPs), and SnpPosition (position in the sequence tag at which the defined SNP variant base occurs) are of particular relevance to our analyses.

### 2.4 SilicoDArT genotyping

We also consider genetic variation as it applies to complementary dominant sequence tag presence-absence markers. These are referred to as SilicoDArT markers, analogous to microarray DArTs first described by Jaccoud *et al*. (2001) but more recently extracted *in silico* from sequences obtained from genomic representations (e.g. Ali *et al*. 2020, Mahboubi *et al*. 2020, Sansaloni, *et al*. 2020, Elshibli and Korpelainen 2021, Nantongo *et al*. 2022). SilicoDArTs are scored in binary (1/0) with score "1" representing presence of restriction fragment while score "0" represents absence of the respective fragment. If the two restriction enzymes (used in the accompanying DArTSeq or ddRAD) find their mark, the corresponding sequence tags are amplified and if the resultant sequences are determined to be homologous, the state is scored as "1" for that locus. An absence "0" can have multiple causes. The most common cause is a mutation (SNP or indel) at one or both of the restriction enzyme sites, whereby the restriction enzyme(s) does not find its mark, and the corresponding sequence tag in that individual is not amplified and sequenced (null allele), or if it is amplified from a different start site, is no longer considered homologous during pre-processing. A second source of a SilicoDArT absence is variation in Cytosine methylation where the restriction enzymes used in complexity reduction are methylation sensitive (Wittenberg *et al*. 2005). A third and smaller source of SilicoDArT absence ("0") is the presence-absence variation (PAVs *sensu* Gabur *et al*. 2020) in the target fragment in the genome. SilicoDArT markers are commonly used in an agricultural setting to generate datasets that are complementary to SNP datasets (Mahboubi *et al*. 2020, Sansaloni, *et al*. 2020, Nantongo *et al*. 2022). They have less commonly been used in the context of population genetics (but see Elshibli and Korpelainen 2021). SilicoDArT markers typically outperform SNP markers in deeper phylogenetics analyses (Alam *et al*. 2018).

Unlike the co-dominant SNP genotype markers, SilicoDArT markers are dominant markers. Technical artefacts such as low DNA quality or quantity, or failure of experimental protocol resulting in shallow sequencing depth, may occasionally lead to missing data which will be erroneously interpreted as a null allele. This is overcome by quality control processes. Specific preprocessing options were used to separate true null alleles (sequence tag absences) from sequence tags present in the target genome but missed by the sequencing because of low read depth. First, the sequence tags were filtered more stringently on read depth than for SNPs, typically with a threshold of 8*x* or more. Second, the distributions of the sequence tag counts were examined for bimodality (clustering on either 0 or 1) and those loci showing a continuum of counts rather than two clusters were discarded. Hemizygous genotypes at the SilicoDArT loci were scored as missing (-). Finally, the scores were tested for reproducibility using technical replicates, providing us the opportunity to filter loci with poor reproducibility in the SilicoDArT scores. In this way, a reliable set of dominant markers are obtained, to complement the SNP dataset generated from the same individuals. Note that although the SilicoDArT and SNP genotypes are derived from the same samples and representation, the SilicoDArT markers also included sequence tags that do not have a SNP, that is, sequence tags that are invariant in their DNA sequence across all individuals.

### 2.5 Additional Filtering

The SNP data and associated metadata were read into a genlight object ({adegenet}, Jombart 2008) to facilitate processing with package dartR (Gruber *et al*. 2018). Standard filters were applied to increase the reliability of the final set of SNP markers. The initial polymorphic SNP loci were filtered to remove all but one SNP per sequence tag, on call rate (threshold 0.95 unless otherwise specified), on repeatability (threshold 0.998) and on read depth (threshold 10*x*). Individuals that had a low call rate across loci (< 0.90) were removed from the analysis (n = 3). Any monomorphic loci arising from the removal of individuals or populations were also deleted. Given the low within-population sample sizes (many n ≤ 10), we did not filter loci for departures from Hardy-Weinberg Equilibrium (HWE) or Linkage Disequilibrium. A companion SilicoDArT dataset for the same individuals was also generated, filtered on read depth (threshold 10*x*), repeatability (threshold 0.998), and call rate (threshold 0.95). We regard the data remaining after this additional filtering of SNP and SilicoDArT genotypes as highly reliable.

### 2.5 Visualization

Genetic SNP similarity among individuals and populations was visualized using ordination (PCA, Jolliffe 2002) as implemented in the gl.pcoa and gl.pcoa.plot functions of dartR. A scree plot of eigenvalues (Cattell, 1966) in combination with the broken-stick criterion (Macarthur, 1957) guided the number of informative axes to examine. While there are alternatives to PCA for visualizing genetic similarities among individuals and populations, we have selected classical PCA because it requires no prior identification of groups or the number of such groups and is assumption free.

### 2.6 Genetic Diversity

Observed and expected heterozygosity (Nei, 1978) and *F_IS_* was obtained for each population from SNP allele frequencies using the gl.report.heterozygosity function of dartR.

### 2.7 Fixed Difference Analysis

A fixed difference between two populations at a locus occurs when the populations share no SNP alleles at that locus. Accumulation of fixed differences between two populations is a robust indication of lack of geneflow. The fixed difference analysis was undertaken using the nuclear SNP data with the gl.fixed.diff and gl.collapse functions in dartR. Populations (that is, field sampling sites) showing evidence of contemporary admixture and hybridization or introgression were identified using NewHybrids (Anderson and Thompson 2002) and eliminated from the analysis. NewHybrids is limited to 200 loci, so we filtered on Average Polymorphic Information Content (AvgPIC) to reduce the number of loci to approximately 200 with minimal loss of relevant information. Nevertheless, because of the stochastic element in the selection of loci implemented in the dartR package, we ran NewHybrids five times on each sample set to check for consistency (after Donnellan *et al*. 2013, Gramlich *et al*. 2018) and averaged the posterior probabilities for class membership (P0, P1, F1, F2, F1xPo, F1xP1) (after Baiakhmetov *et al*. 2021). Populations with low sample sizes (< 3) were amalgamated with other populations from the same or adjacent drainages before analysis – those within each of South Alligator River (n=1), Reynolds River (n=1), Towns River (n=1), Darwin River (n=1) and McKinlay River (n=2). Fixed differences were counted for populations taken pairwise, and when two populations had no fixed differences or only one fixed difference (i.e. lacked corroboration), they were combined and the process repeated until there was no further reduction (Georges and Adams, 1996). The resultant entities are, by definition, putatively diagnosable at two or more SNP loci.

The decision to amalgamate two populations can be made with certainty because lack of fixed allelic differences at all loci in the samples at hand cannot be eroded by the addition of more data. However, the separation of two populations by one or more fixed differences is subject to sampling error. False positives may arise because of the finite sample sizes involved. Simulations as implemented in dartR were used to estimate the expected false positive rate in pairwise comparisons (Georges *et al*. 2018). Populations or diagnosable aggregations with an observed count of fixed differences not significantly different from the expected rate of false positives were further amalgamated to yield a final set of diagnosable aggregations.

### 2.9 Phylogeny

We estimated the phylogeny from the nuclear SNP data using SVDquartets analysis (Chifman and Kubatko 2014, 2015), chosen because of our short reads (≤ 69 bp) and the typically single variable sites per locus (Chou *et al*. 2015). Heterozygous SNP positions were represented by standard ambiguity codes (see Felsenstein 2004:255). SVDquartets takes unlinked multi-locus data for subsets of taxa, taken four at a time (quartets), and assigns a score to each of the three possible unrooted topologies for each quartet. The topology with the lowest score is selected as the best supported topology for that quartet. The final set of quartets is combined (Reaz *et al*. 2014) to estimate the species tree. We used the implementation of SVDquartets in PAUP* (version 4.0a169; Swofford 2003) with parameters evalQuartets=random, bootstrap=standard, nreps=10,000, ambigs=distribute. The tree was provisionally rooted with the southern *Emydura*.

We estimated the phylogeny from the SilicoDArT data (presence-absence of sequence tags) using parsimony as implemented in PAUP*. Because of prohibitive computational time, the analysis was subset to 1000 jobs run concurrently, each corresponding to one bootstrap replicate, on the Gadi computer array of the Australian National Computational Infrastructure (NCI, https://nci.org.au/) with parameters bootstrap nreps=1 search=heuristic / start=stepwise addseq=random nreps=100 swap=TBR. The resultant bootstrap replicates were combined using PAUP contree all / strict=no majrule=yes percent=50. Again, the tree was provisionally rooted with the southern *Emydura*.

Phylogenetic analyses of whole mitochondrial sequences were performed with maximum likelihood (ML) using the IQTREE web implementation (Trifinopoulos *et al*. 2016, http://iqtree.cibiv.univie.ac.at, accessed 16-Mar-25; Nguyen *et al*. 2015) with parameters -m TESTNEW -bb 1000 -alrt 1000. Modelfinder compared 286 substitution models and selected the General Time Reversable model GTR+F+R2 as best against all criteria. Phylogenetic analysis of the *cytB* sequences were similarly run on IQTREE with parameters -m GTR+R4+F -bb 1000 -alrt 1000. Details of specimens used for the *cytB* sequencing and phylogenetic analysis are shown in Table S2 and for whole mitochondrial sequencing and phylogenetic analysis are shown in Table S3.

## 3 RESULTS

### 3.1 SNP Dataset

#### Step 1: Comprehensive sampling

A total of 100,848 polymorphic SNP loci were scored for 526 individuals of *Emydura* from 43 drainage basins of northern Australia and Papua New Guinea (Figure 2). A total of 39,943 secondary SNPs (additional SNPs sharing the same sequence tag, sensu Gruber *et al*. 2018) were filtered to yield independent 60,905 SNPs. After stringent filtering on repeatability (repAvg < 0.998) calculated from technical replicates and call rate (< 0.95), the number of SNP loci in the dataset dropped to 20,346 and then 13,102 respectively. Three specimens had a call rate of less than 90% (UC_1509[Emtan_Daly], EG_EW17[Emwor_Roper], UC_1549[Emaus_Daly]) and were removed from the dataset. Monomorphic loci (n = 16) arising from the removal of these individuals were deleted. Finally, an additional 1,514 SNPs were filtered because their sequence tags had a read depth less than 10*x*. The resultant dataset comprised 11,572 polymorphic SNP loci from 523 individuals from 43 populations (drainage basins) (n = 1–63).

The corresponding SilicoDArT dataset comprised 132,890 loci scored for the 526 individuals from 43 populations. After filtering for Callrate (< 0.95), Repeatability (< 0.998) and read depth (< 10*x*), there were 73,220 loci scored.

### 3.2 Qualitative Analysis

#### Step 2: Removal of individuals subject to putative hybridization

Preliminary analysis of the data using NewHybrids (Anderson and Thompson 2002) applied to the species taken pairwise revealed clear evidence of a low incidence of admixture arising from contemporary hybridization (Table 1). There are three primary foci for admixture between species. The Daly River has 14 individuals showing evidence of contemporary admixture between *E. tanybaraga* and *E. australis*, out of the 241 animals examined. Seven of these were difficult to assign to species at the time of collection indicating the admixture was also evident morphologically. There was also one individual putatively admixed between *E. tanybaraga* and *E. australis* in the Reynolds River. Two additional specimens showed minor signs of admixture and were conservatively removed also. All 5 individuals in the Flinders River showed evidence of admixture between *E. tanybaraga* and *E. subglobosa worrelli* and were also difficult to assign to species at the time of collection. In addition, there was one putatively admixed individual in the Nicholson River and two in the Roper River, one of which was a putative F1 hybrid. These were assigned to *E. subglobosa worrelli* at time of collection. A third focus for admixture was the Barron and Russell-Mulgrave rivers of east coastal Queensland, as was flagged but not resolved by Georges *et al*. (2018). Evidence from the present study indicates the likely dispersal of *E. tanybaraga* from the Mitchell River (Q) into Tinaroo Dam on the Barron River above the escarpment leading to hybridization and admixture with *E. macquarii* [NthQld]. There is evidence of further dispersal of *E. tanybaraga* or admixed individuals to the lower Barron River and the Russell-Mulgrave River (Table 1).

**Table 1.** Evidence of contemporary hybridization and admixture among the five currently recognised taxa *Emydura australis*, *E. subglobosa subglobosa*, *E. subglobosa worrelli*, *E. tanybaraga* and *E. macquarii* [NthQLD]. Field identities are the taxon names assigned to the individuals on capture -- those marked as [yellowface] were regarded as of ambiguous identity. Values in the body of the table for the parentals are counts of animals assigned concordantly with their field identification. Values in the body of the table for the listed individuals are the posterior probabilities of membership to the assignment categories of NewHybrids (Anderson and Thompson 2002), averaged over 5 runs (Table S4), for those individuals for which assignment to parental taxa was ambiguous. These are likely admixed individuals. EG_EW03 from the Roper River was identified during field collection as *E. s. worrelli* but is likely an F1 hybrid between *E. s. worrelli* and *E. tanybaraga*. Numbers of loci in square brackets after the comparison titles are the number of loci retained after filtering on Polymorphic Information Content (AvgPIC) to reduce loci to ca 200 (a limitation of NewHybrids) with minimal loss of information.

| Taxon/ID | Field Identity | P0 | P1 | F1 | F2 | F1xP0 | F1xP1 | CytB HaploClade |
| --- | --- | --- | --- | --- | --- | --- | --- | --- |
| ***E. tanybaraga* vs *E. australis* [217 loci]** |  |  |  |  |  |  |  |  |
| P0:Emtan | *E. tanybaraga* | 73 | 0 | 0 | 0 | 0 | 0 |  |
| P1:Emaus | *E. australis* | 0 | 154 | 0 | 0 | 0 | 0 |  |
| AA072622:Daly | *E. australis* | 0.00 | 0.47 | 0.00 | 0.50 | 0.00 | 0.03 | -- |
| AA072645:Daly | *E. australis* | 0.00 | 0.24 | 0.00 | 0.37 | 0.00 | 0.39 | -- |
| UC_0269:Daly | *E. australis* | 0.00 | 0.31 | 0.00 | 0.34 | 0.00 | 0.35 | Hap38_Emaus_Daly(35) |
| UC_0383:Daly | *E. australis* | 0.00 | 0.00 | 0.00 | 0.99 | 0.00 | 0.01 | Hap38_Emaus_Daly(35) |
| UC_0386:Daly | *E. australis* | 0.00 | 0.61 | 0.00 | 0.10 | 0.00 | 0.29 | Hap38_Emaus_Daly(35) |
| UC_0841:Daly | *E. [yellowface]* | 0.00 | 0.00 | 0.00 | 1.00 | 0.00 | 0.00 | -- |
| UC_0929:Daly | *E. tanybaraga* | 0.00 | 0.00 | 0.00 | 0.98 | 0.01 | 0.01 | Hap38_Emaus_Daly(35) |
| UC_1508:Daly | *E. [yellowface]* | 0.21 | 0.00 | 0.00 | 0.67 | 0.12 | 0.00 | Hap39_Emaus_Daly(5) |
| UC_1514:Daly | *E. [yellowface]* | 0.02 | 0.00 | 0.00 | 0.75 | 0.23 | 0.00 | Hap38_Emaus_Daly(35) |
| UC_1517:Daly | *E. [yellowface]* | 0.79 | 0.00 | 0.00 | 0.14 | 0.07 | 0.00 | Hap39_Emaus_Daly(5) |
| UC_1519:Daly | *E. tanybaraga* | 0.00 | 0.00 | 0.00 | 0.73 | 0.26 | 0.00 | Hap37_Emtan_Daly(1) |
| UC_1521:Daly | *E. [yellowface]* | 0.00 | 0.00 | 0.00 | 0.99 | 0.00 | 0.01 | Hap38_Emaus_Daly(35) |
| UC_1536:Daly | *E. [yellowface]* | 0.00 | 0.00 | 0.00 | 0.96 | 0.04 | 0.00 | Hap38_Emaus_Daly(35) |
| yface93:Daly | *E. [yellowface]* | 0.79 | 0.00 | 0.00 | 0.15 | 0.06 | 0.00 | Hap40_Emtan_Daly(1) |
| UC_0661:Reynolds | *E. tanybaraga* | 0.00 | 0.00 | 0.00 | 0.78 | 0.22 | 0.00 | Hap29_Emtan_Reyn(1) |
| ***E. tanybaraga* vs *E. s. worrelli* [621 loci]** |  |  |  |  |  |  |  |  |
| P0:Emtan | *E. tanybaraga* | 73 | 0 | 0 | 0 | 0 | 0 |  |
| P1:Emwor | *E. s. worrelli* | 0 | 125 | 0 | 0 | 0 | 0 |  |
| HBS_32227:Nicholson | *E. s. worrelli* | 0.00 | 0.58 | 0.00 | 0.12 | 0.00 | 0.31 | -- |
| EG_EW03:Roper | *E. s. worrelli* | 0.04 | 0.01 | 0.80 | 0.09 | 0.02 | 0.04 | -- |
| HBS_32131:Roper | *E. s. worrelli* | 0.00 | 0.79 | 0.00 | 0.06 | 0.00 | 0.15 | -- |
| AA046038:Flinders | *E. [yellowface]* | 0.40 | 0.20 | 0.00 | 0.40 | 0.00 | 0.00 | -- |
| AA046039:Flinders | *E. [yellowface]* | 0.20 | 0.20 | 0.00 | 0.58 | 0.00 | 0.02 | -- |
| AA046040:Flinders | *E. [yellowface]* | 0.20 | 0.20 | 0.00 | 0.55 | 0.00 | 0.05 | -- |
| AA046041:Flinders | *E. [yellowface]* | 0.20 | 0.20 | 0.00 | 0.60 | 0.00 | 0.00 | -- |
| AA046042:Flinders | *E. [yellowface]* | 0.20 | 0.20 | 0.00 | 0.55 | 0.05 | 0.00 | -- |
| ***E. tanybaraga* vs *E. s. subglobosa* [264 loci]** |  |  |  |  |  |  |  |  |
| P0:Emtan | *E. tanybaraga* | 73 | 0 | 0 | 0 | 0 | 0 |  |
| P1:Emsub | *E. s. subglobosa* | 0 | 147 | 0 | 0 | 0 | 0 |  |
| ***E. s. subglobosa* vs *E. s. worrelli* [205 loci]** |  |  |  |  |  |  |  |  |
| P0:Emsub | *E. s. subglobosa* | 147 | 0 | 0 | 0 | 0 | 0 |  |
| P1:Emwor | *E. s. worrelli* | 0 | 125 | 0 | 0 | 0 | 0 |  |
| ***E. s. worrelli* vs *E. australis* [228 loci]** |  |  |  |  |  |  |  |  |
| P0:Emwor | *E. s. worrelli* | 0 | 125 | 0 | 0 | 0 | 0 |  |
| P1:Emaus | *E. australis* | 153 | 0 | 0 | 0 | 0 | 0 |  |
| ***E. s. subglobosa* vs *E. australis* [212 loci]** |  |  |  |  |  |  |  |  |
| P0:Emsub | *E. s. subglobosa* | 0 | 147 | 0 | 0 | 0 | 0 |  |
| P1:Emaus | *E. australis* | 153 | 0 | 0 | 0 | 0 | 0 |  |
| ***E. macquarii* [NthQLD] vs *E. tanybaraga* [Mitchell(Q)] [396 loci]** |  |  |  |  |  |  |  |  |
| P0:Emmac | *E. macquarii* | 69 | 0 | 0 | 0 | 0 | 0 |  |
| P1:Emtan | *E. tanybaraga* | 0 | 13 | 0 | 0 | 0 | 0 |  |
| 98EmCairns01:Barron | *E. macquarii* | 0.00 | 0.00 | 0.00 | 0.00 | 1.00 | 0.00 | -- |
| 98EmCairns05:Barron | *E. macquarii* | 0.00 | 0.00 | 0.12 | 0.88 | 0.00 | 0.00 | -- |
| 98EmCairns07:Barron | *E. macquarii* | 0.00 | 0.00 | 0.00 | 0.60 | 0.40 | 0.00 | -- |
| 98EmCairns08:Barron | *E. macquarii* | 0.20 | 0.00 | 0.00 | 0.00 | 0.80 | 0.00 | -- |
| AA037115:Barron [Tinaroo] | *E. [yellowface]* | 0.00 | 0.00 | 0.00 | 0.00 | 1.00 | 0.00 | Hap36_Emtan_MitQ-Holr-Staa-Wentl-Arch-Pasc-Barr(17) |
| AA037116:Barron [Tinaroo] | *E. [yellowface]* | 0.00 | 0.00 | 0.00 | 1.00 | 0.00 | 0.00 | Hap36_Emtan_MitQ-Holr-Staa-Wentl-Arch-Pasc-Barr(17) |
| HBS_101659:Mulgrave-Russell | *E. macquarii* | 0.00 | 0.00 | 1.00 | 0.00 | 0.00 | 0.00 | -- |
| HBS_101677:Mulgrave-Russell | *E. macquarii* | 0.00 | 0.00 | 1.00 | 0.00 | 0.00 | 0.00 | -- |

As contemporary hybridization is problematic for both the fixed difference analyses and the phylogenetic analyses, these putative hybrid and backcrossed individuals from the northern rivers were conservatively removed from subsequent analyses. The dataset was, as a consequence, reduced to 498 individuals from 41 populations scored for 10,445 SNPs. For phylogenetic analysis using nuclear data, we added 10 individuals of *Emydura macquarii* from the Murray-Darling Basin, 10 from the Lake Eyre Basin and 9 from the Fitzroy River of Queensland) drawing from the dataset of Georges *et al*. (2018). In this full dataset, including the 29 additional individuals from *E. macquarii*, 12,420 SNP loci were polymorphic for the 527 individuals from 44 drainage basins. The corresponding SilicoDArT dataset was polymorphic for 45,945 loci from 528 individuals from 44 populations. For the mitochondrial data, we added sequence from one specimen of *Elseya flaviventralis* to serve as outgroup taxon.

#### Step 3: Exploratory analysis based on PCA

A PCA generated from the SNP data after removal of putative admixed individuals yielded 8 informative dimensions (broken-stick criterion, Macarthur 1957, Jackson 1993) from 513 original dimensions. The top three dimensions explained 58.1% of the total variance. Four distinct groupings were evident in the PCA corresponding to aggregations of populations from each of the four currently recognised taxa *Emydura australis, E. tanybaraga, E. subglobosa subglobosa* and *E. subglobosa worrelli* (Figure 3). The distinction between *E. subglobosa subglobosa* and *E. subglobosa worrelli* was strongly evident in the third dimension of the PCA (12.6% of variance explained). The population of *Emydura* in the Jardine River of Cape York Queensland grouped with the populations of *E. subglobosa subglobosa* from Papua New Guinea. This confirms its species identity as *E. subglobosa* (Cogger 2018) formerly established on the basis of morphology and coloration. Genetic Euclidean distances between populations of the currently-recognised taxa averaged 32.5±3.88 (11.4-37.3, n=456) compared with distances between populations within the same taxon (14.3±3.15 (6.7-23.5, n=139) (Table 2).

**Figure 3.**
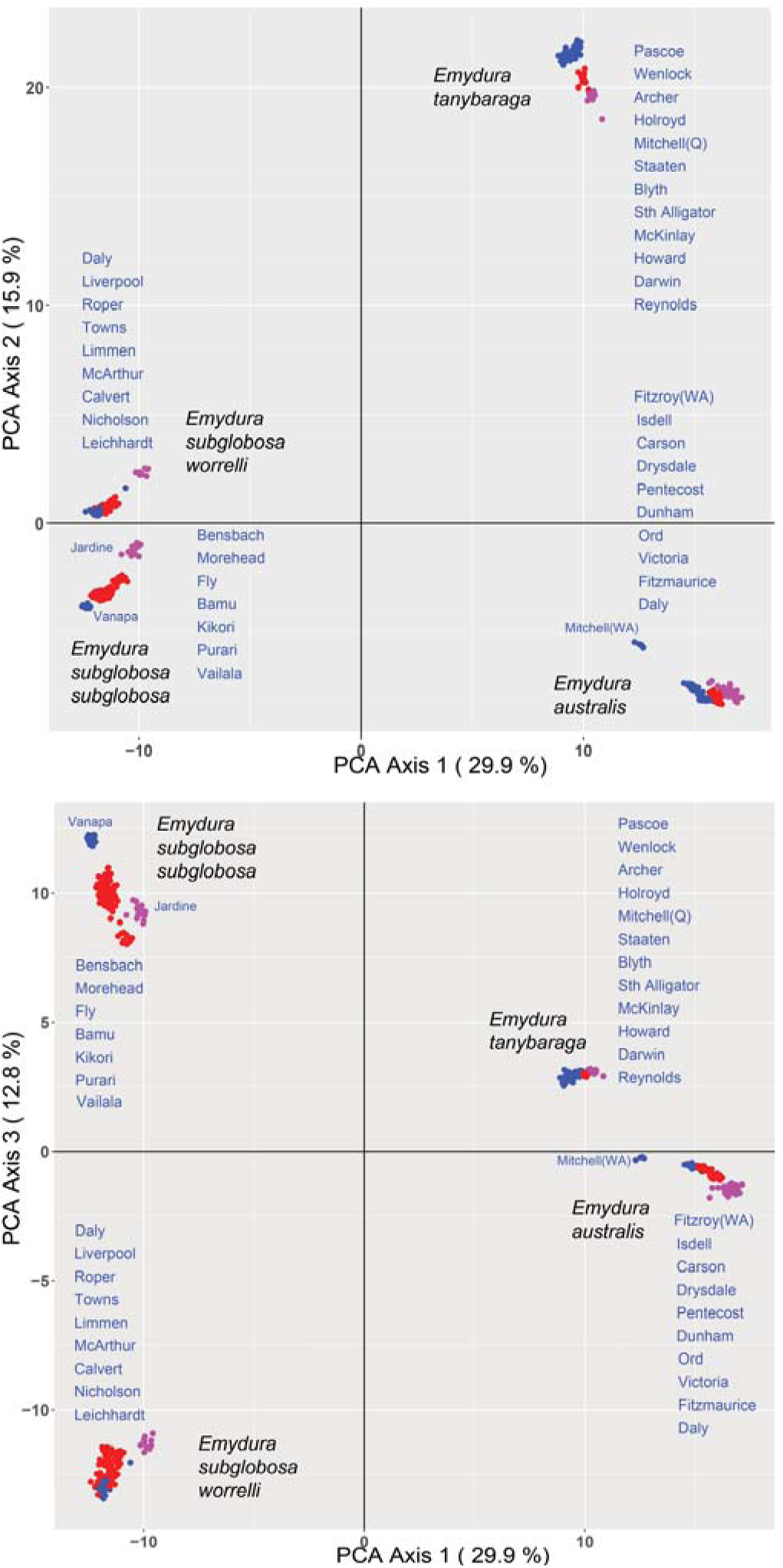
A principal components analysis (PCA) with populations (sampling localities) as entities, SNP loci as attributes and the SNP scores (0, 1, 2) as character states. There are four major groupings. River drainages from which the specimens were collected are listed for each major group. Colours are only to facilitate cross comparison between the plots for Axis 1 and 2 versus Axis 1 and 3. Populations from the Mitchell River (WA), the Jardine River and the Vanapa River are labelled because they are mentioned specifically in the text.

**Table 2.** Euclidean genetic distances between major groupings of the PCA averaged across their populations, and averaged for populations within each of the major groupings of the PCA. For a full distance matrix, refer to Table S6.

| Taxa |  | Mean | Min | Max | SD | N |
| --- | --- | --- | --- | --- | --- | --- |
| Emaus | Emsub | 33.8 | 31.8 | 37.5 | 1.34 | 99 |
|  | Emtan | 33.2 | 31.0 | 34.7 | 0.85 | 88 |
|  | Emwor | 34.3 | 33.0 | 35.8 | 0.66 | 88 |
| Emsub | Emtan | 35.7 | 33.8 | 39.2 | 1.32 | 72 |
|  | Emwor | 26.7 | 24.0 | 31.0 | 1.75 | 72 |
| Emtan | Emwor | 35.5 | 33.3 | 36.8 | 0.81 | 64 |
| Overall |  | 33.2 | 24.0 | 39.2 | 1.17 | 483 |
|  | Emaus | 16.0 | 6.7 | 23.5 | 4.08 | 55 |
|  | Emwor | 13.2 | 9.2 | 15.9 | 1.96 | 28 |
|  | Emsub | 14.6 | 7.0 | 22.9 | 3.86 | 36 |
|  | Emtan | 12.6 | 7.4 | 15.4 | 1.90 | 28 |
| Overall |  | 14.5 | 6.7 | 23.5 | 3.33 | 147 |

As there were 8 informative dimensions in the PCA, there is additional information not represented in the plots of Axes 1-3, revealed in PCA subplots generated for each of the four major groupings (after Georges and Adams 1992, Unmack *et al*. 2022) (Figure 4). *Emydura australis* was represented by three groupings in the PCA based on divergent allelic frequencies. A western Kimberley grouping (Fitzroy River to the Drysdale River) and an eastern Kimberley grouping (Ord River to the Fitzmaurice River) are presumably isolated by the eastern plateau of the Kimberley (Karunjie Plateau). The plateau is deeply dissected by the Drysdale River in the west and the rivers draining into the Ord River estuary in the east, such that their headwaters come into close proximity, but the waters of the gorges in this deeply dissected landscape is unsuitable habitat for turtles and the steep terrain not suitable for turtle dispersal across catchment boundaries. A third grouping of *E. australis* occurs in the Daly River.

**Figure 4.**
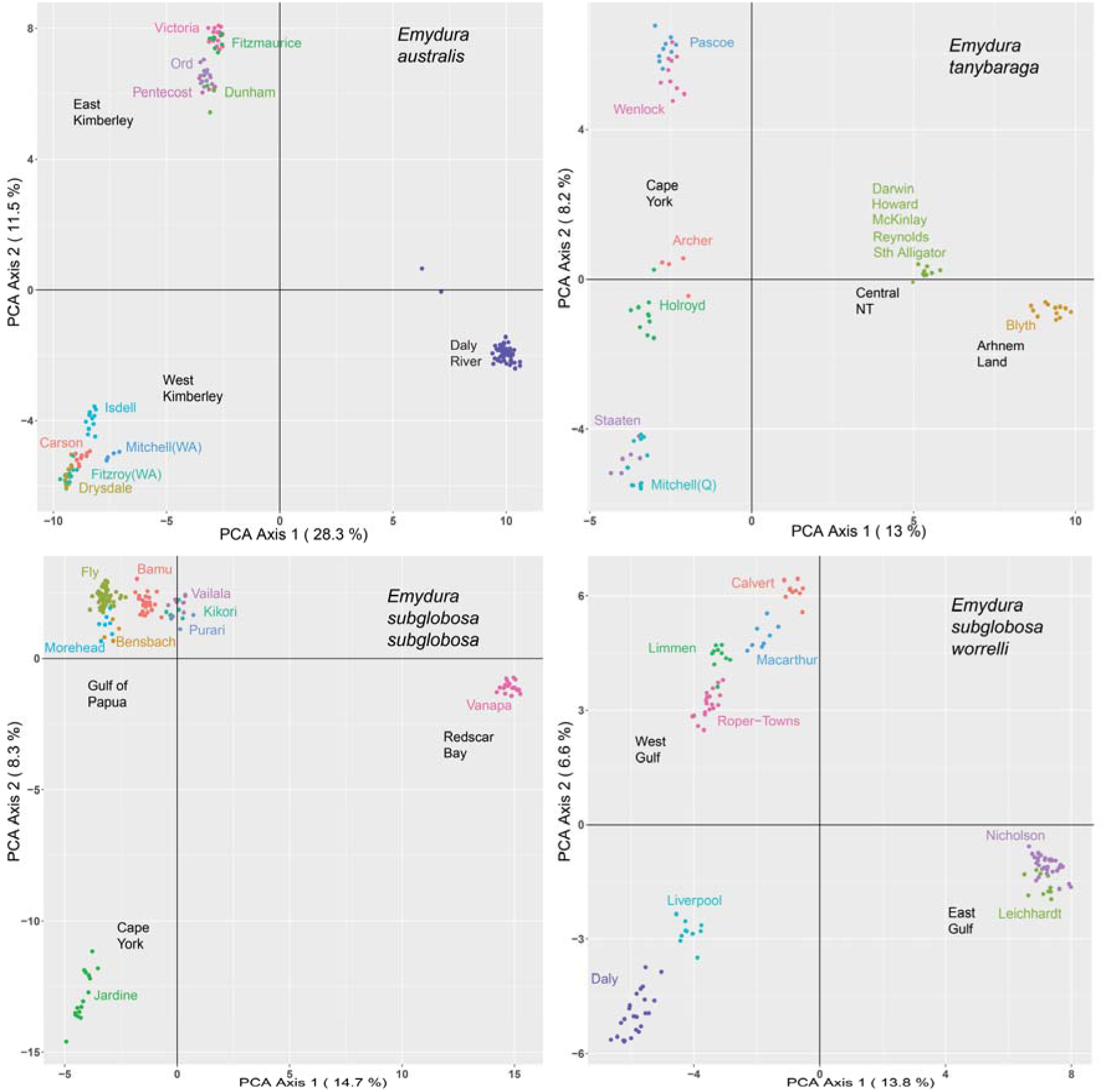
Supplementary plots for PCA applied to each of the four major groupings in the primary PCA plot.

*Emydura tanybaraga* has a disjunct distribution, occurring as geographically separated populations in Queensland and in the Northern Territory (Georges and Adams 1996, Georges and Thomson 2010), confirmed by our current analysis. The Queensland populations form a series, faithful to physical geography, along the west flowing rivers of Cape York Peninsula, from the Mitchell River in the south to the Wenlock River in the north where it crosses to the adjacent eastern flowing Pascoe River (Figure 4). In the Northern Territory, the species forms two groupings, one in the Darwin region (east to the South Alligator River) and one in Arnhem Land (Blyth River). *Emydura tanybaraga* occurs also in the Daly River of the Northern Territory and the Flinders River of Queensland, but all sampled individuals showed evidence of introgression with *E. australis* and *E. s. worrelli* respectively (Table 1); they were excluded from subsequent analysis including the PCA subplots of Figure 4.

*Emydura subglobosa subglobosa* is primarily distributed in southern New Guinea in the rivers flowing south from the central ranges. Samples from Indonesian New Guinea were not available to us. The taxon in Papua New Guinea formed a major grouping corresponding to populations in the rivers flowing south into the Gulf of Papua, from the Bensback River in the west to the Vailala River in the east (Figure 2). With the exception of the Vailala River, these rivers are interconnected in their lowlands by extensive freshwater swamps, mangroves and associated channels, providing avenues for dispersal of *Emydura*. Nevertheless, the PCA shows a series of populations extending across the gulf that are broadly respectful of physical geography. A second grouping in Papua New Guinea corresponds to populations in the Vanapa River region (including the Laloki, Martin, Vakabu and Vanapa rivers). A third grouping corresponds to the population in the small Jardine River at the tip of Cape York Peninsula in Australia.

*Emydura subglobosa worrelli* is distributed in the rivers discharging into the Gulf of Carpentaria from the Roper River in the west to the Flinders in the east, again represented by a series of populations that reflects physical geographic proximity. All the animals sampled from the Flinders River showed evidence of introgression with *E. tanybaraga* (Table 1) and so are not shown in Figure 4. The taxon extends westward into the rivers draining Arnhem Land to the north, including the Liverpool and Daly drainages where it primarily occupies the plateau regions above the Arnhem Land escarpment. *E. tanybaraga* and *E. australis* occupy the middle and lowland reaches below the escarpment.

It is important to note that the groupings in the PCAs represent differences in allele frequency profiles and not necessarily differences that are diagnostic at the individual level. Diagnostic differences are revealed by a fixed difference analysis.

#### Step 4. Fixed difference analysis

We examined populations (collection sites) pairwise for fixed allelic differences using the gl.fixed.diff and gl.collapse functions in dartR and amalgamated populations for which there were no corroborated fixed differences (nloc=1). This analysis assumed no *a priori* assignment to species. Such an analysis is highly sensitive to admixture and, in particular, to the presence of F1 hybrids, hence the removal of any individuals showing evidence of putative admixture in the NewHybrids analysis. It is also highly sensitive to false positives arising from low sample sizes, so where possible, we manually amalgamated populations for which we had only one or two individuals with an appropriate neighbour. For example, we had only one individual from the Towns River. This river shares a floodplain with the adjacent Roper River, so we amalgamated the single individual from the Towns River with the 20 individuals from the Roper River manually, before the fixed difference analysis. The one individual from the Darwin River, 2 individuals from the McKinlay River and 2 individuals from the Reynolds River were amalgamated manually with the 5 individuals from the Howard River to form a Central NT grouping. To this aggregation, we manually added the one individual from the South Alligator River based on its proximity in the PCA (Figure 4).

The fixed difference analysis yielded five diagnosable aggregations. Four of these corresponded the currently recognised taxa, and the counts of fixed differences in each case significantly exceeded the false positive rate (Table 3, upper triangle). A fifth diagnosable aggregation corresponded to the population of *E. australis* in the Kimberley’s Mitchell River above the escarpment (Table 3). The count of fixed differences between this Mitchell River population and remaining populations of *E. australis* marginally exceeded the false positive rate (p = 0.491) at the 5% level of significance. These five diagnosable aggregations are consistent with the groupings evident in the first two axes of the PCA. The fixed difference analysis provided independent confirmation of the currently-recognised four taxa. They are candidates for species level classification, subject to additional phylogenetic consideration.

**Table 3.** A matrix of the counts of fixed allelic differences (lower triangle) between diagnosable entities identified using a fixed difference analysis. All were highly significant (p < 0.0001) (p values above diagonal) bar the comparison between *E. australis* from the Mitchell River and remaining populations of *E. australis*, which was marginally significant (p = 0.0491).

|  | *E. australis* | *E. australis* [Mitchell] | *E. s. subglobosa* | *E. tanybaraga* | *E. s. worrelli* |
| --- | --- | --- | --- | --- | --- |
| *E. australis* |  | 0.0491 | 0.0000 | 0.0000 | 0.0000 |
| *E. australis* [Mitchell] | 23 |  | 0.0000 | 0.0000 | 0.0000 |
| *E. s. subglobosa* | 53 | 302 |  | 0.0000 | 0.0000 |
| *E. tanybaraga* | 94 | 354 | 121 |  | 0.0000 |
| *E. s. worrelli* | 51 | 289 | 7 | 84 |  |

#### Step 5. Phylogeny

The SVDquartets tree constructed from the SNP genotypes generated a tree with strong bootstrap support across many nodes (Figure 5a). Each of the populations assigned to diagnosable aggregations in the fixed difference analysis corresponded to major lineages in the phylogeny, namely *Emydura australis*, *E. tanybaraga*, *E. subglobosa subglobosa* and *E. subglobosa worrelli* each with strong bootstrap support. Shallower lineages with strong bootstrap support (> 90%) included a lineage of *E. australis* in the Daly River distinct from the remaining populations of this species. There was also a distinction between populations draining the western Kimberley Plateau (Fitzroy, Drysdale, Carson, Isdell) from the eastern Kimberley Plateau (Ord, Pentecost, Dunham, Fitzmaurice, Victoria), with the Mitchell River population joining these two in a polytomy. There was a distinction between the Queensland and Northern Territory lineages of *E. tanybaraga*; lineage structure among the populations of *E. subglobosa subglobosa* of Papua New Guinea; and a distinction with 100% bootstrap support between the Jardine River population of *Emydura s. subglobosa* and its Papua New Guinea counterparts. There was also lineage structure evident among the populations of *E. worrelli*. None of these shallower lineages corresponded to uniquely diagnosable aggregations notwithstanding their high bootstrap support.

**Figure 5.**
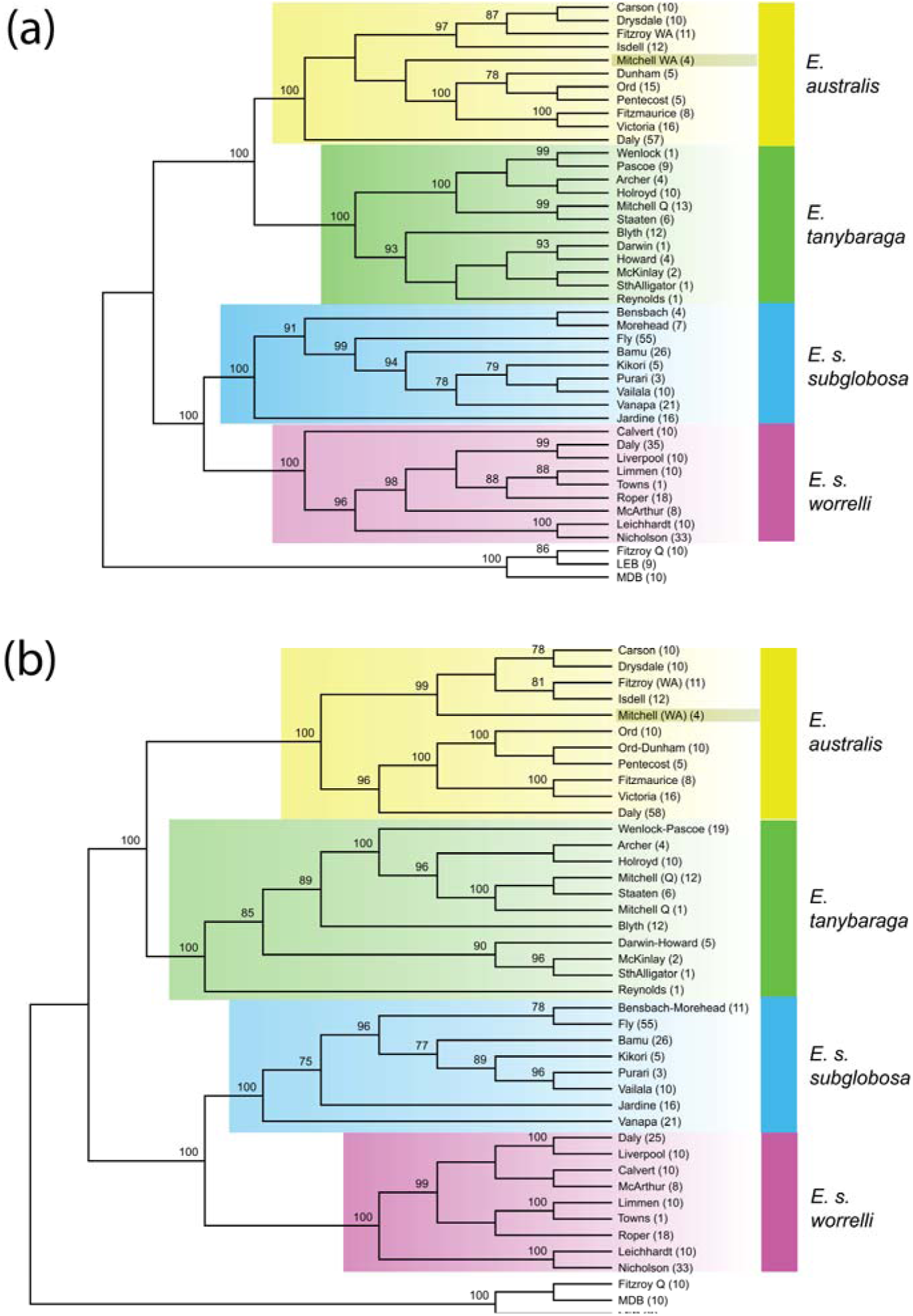
Phylogenies for the populations of *Emydura australis, E. tanybaraga, E. subglobosa subglobosa* and *E. s. worrelli* based on (a) SVDquartets applied to SNP genotypes with ambiguity codes substituted for heterozygous sites and (b) parsimony analysis applied to SilicoDArT presence-absence markers. Trees are rooted with the southern Emydura (Fitzroy River, Qld, Murray-Darling Basin (MDB), Lake Eyre Basin (LEB) in the absence of an alternative sufficiently close outgroup taxon suitable for SNP analysis. Bootstrap values are shown on nodes; they are based on 100 replicates for the SVDquartets and 1000 replicates for the parsimony analysis. The coloured bars identify diagnosable lineages arising from the fixed allelic difference analysis. They are concordant with the contemporary taxa at species and subspecies level. The enigmatic Mitchell River population from the Kimberley is as marked. Refer to Table S1 for specimens examined.

The parsimony analysis applied to the SilicoDArT presence-absence data (Figure 5b) yielded comparable results to those outlined above for the SVDquartets analysis with slightly better resolution and bootstrap support for shallower lineages. Again, the diagnosable aggregations identified in the fixed difference analysis corresponded well to major clades that could be assigned to currently recognised taxa.

The maximum likelihood (ML) phylogeny generated for mitochondrial *cytB* drew from 1,108 bases generated from 57 distinct haplotypes. Of these, 865 bases were constant across haplotypes, 183 were variable but parsimony uninformative and 160 were parsimony informative. ML recovered one tree with an ln score of -3374.80 ± 107.31SE (Figure 6). Three major clades corresponded with the currently recognised taxa *Emydura australis*, *E. subglobosa subglobosa* and *E. s. worrelli*. The clade that contained *E. tanybaraga* was less well defined. Surprisingly, the mitochondrial haplotype of *E. s. subglobosa* from the Jardine River showed close relationship with the clade associated with *E. tanybaraga*. The Daly River individuals of *E. australis* showed evidence of admixture with *E. tanybaraga* (Table 1; red dots in Figure 6) and, apart from one admixed individual (UC_1519, haplotype 78 of Figure 6), fell internal to the clade that included *E. tanybaraga*. The Kimberley’s Mitchell River population was again distinctive. It comprised two clades, one contributing to an *E. australis* clade (haplotypes 88, 90, 91) consistent with its placement in the nuclear marker phylogenies; the other fell within the clade that that included *E. tanybaraga* (haplotype 25, n = 7). In these two respects, the relationships of *E. s. subglobosa* from the Jardine River and *E. australis* from the Mitchell River (WA), the mitochondrial phylogeny departed in substantial ways from the nuclear phylogeny.

**Figure 6.**
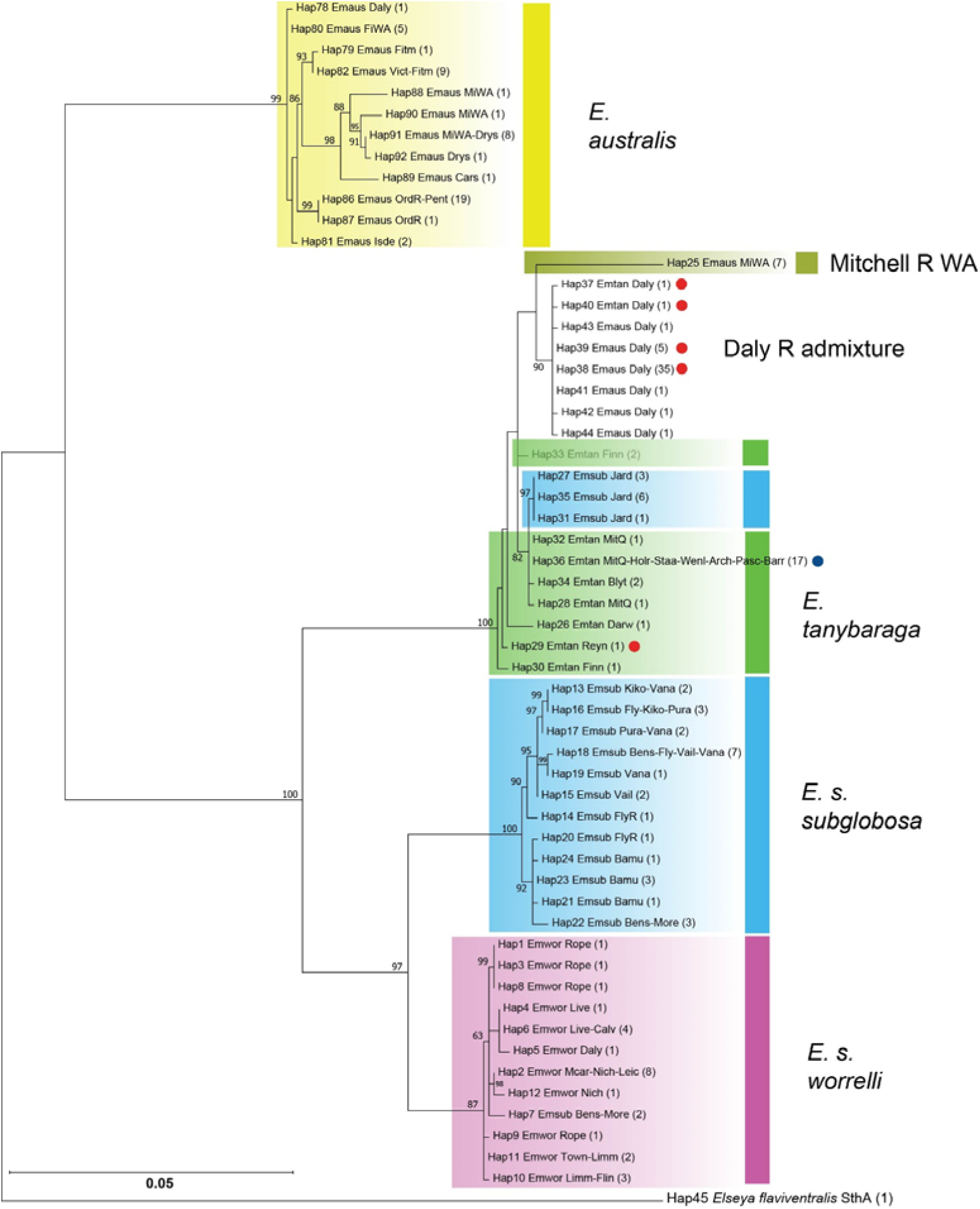
Maximum likelihood phylogeny for the northern *Emydura* based on 1109 bp of Cytochrome B mtDNA sequence. Taxon labels include the haplotype number, the five letter species abbreviation, the four letter drainage basin abbreviation(s) and the number of sequences with the haplotype. Table S5 gives the individuals assigned to each haplotype. The red dots show haplotypes that occur in some individuals showing evidence of admixture between *Emydura tanybaraga* and *Emydura australis* (Table 1). The blue dot refers to populations of *Emydura tanybaraga* showing evidence of admixture with *Emydura macquarii krefftii* in the east coastal Barron River (Tinaroo Dam). Coloured bars refer to the diagnosable entities identified in the fixed difference analysis. The Daly River shows a high level of admixture (Table 1). Bootstraps are based on 1000 replicates, only those with strong bootstrap support (> 80%) are shown. Refer to Table S1 for specimens examined.

The phylogeny generated from whole mtDNA sequences (Figure 7) had strong bootstrap support for all nodes but one, and reinforced evidence for the sister status of *E. subglobosa subglobosa* (including the holotype LR215683, Kehlmaier *et al*. 2019) and *E. s. worrelli* (Georges and Adams 1992). There was strong support also for an *Emydura australis* clade consistent with the diagnosable aggregations defined by the fixed difference analysis. This clade contained the holotypes for *Emydura australis* (LR215684, Kehlmaier *et al*. 2019) and *Emydura victoriae* (LR215686, Kehlmaier *et al*. 2019) which is consistent with the aggregation of *E. australis* including those from the type locality of *E. victoriae* in the fixed difference analysis. The sister relationship between *E. australis* and *E. tanybaraga* (Georges and Adams 1992) was not supported because the southern *Emydura macquarii* was internal to the clade representing the northern *Emydura*. The Mitchell River (WA) haplotypes (haploclade 1, n=6) fall into the *Emydura australis* clade whereas other Mitchell River haplotypes (haploclade 2, n=5) fell within the clade containing the *Emydura tanybaraga* haplotype suggesting the presence of both species in the Mitchell River (WA). The haplotype of Daly River specimen UC_0245 (KY857554) fell within the clade containing *Emydura tanybaraga* consistent evidence of admixture between *E. australis* and *E. tanybaraga* in the Daly River (Table 1); it had the characteristic expanded mouth plate of *E. australis* (used in the field to assign it to *E. australis*) but an *E. tanybaraga* mtDNA haplotype. The whole mitochondrial analysis (Figure 7) reinforced the finding from the *cytB* analysis (Figure 6) that *E. s. subglobosa* from the Jardine River at the tip of Cape York in Australia bore close relationship with the mt genome sequence of *E. tanybaraga* rather than the clade for *E. s. subglobosa* from New Guinea that was expected from the nuclear DNA analysis.

**Figure 7.**
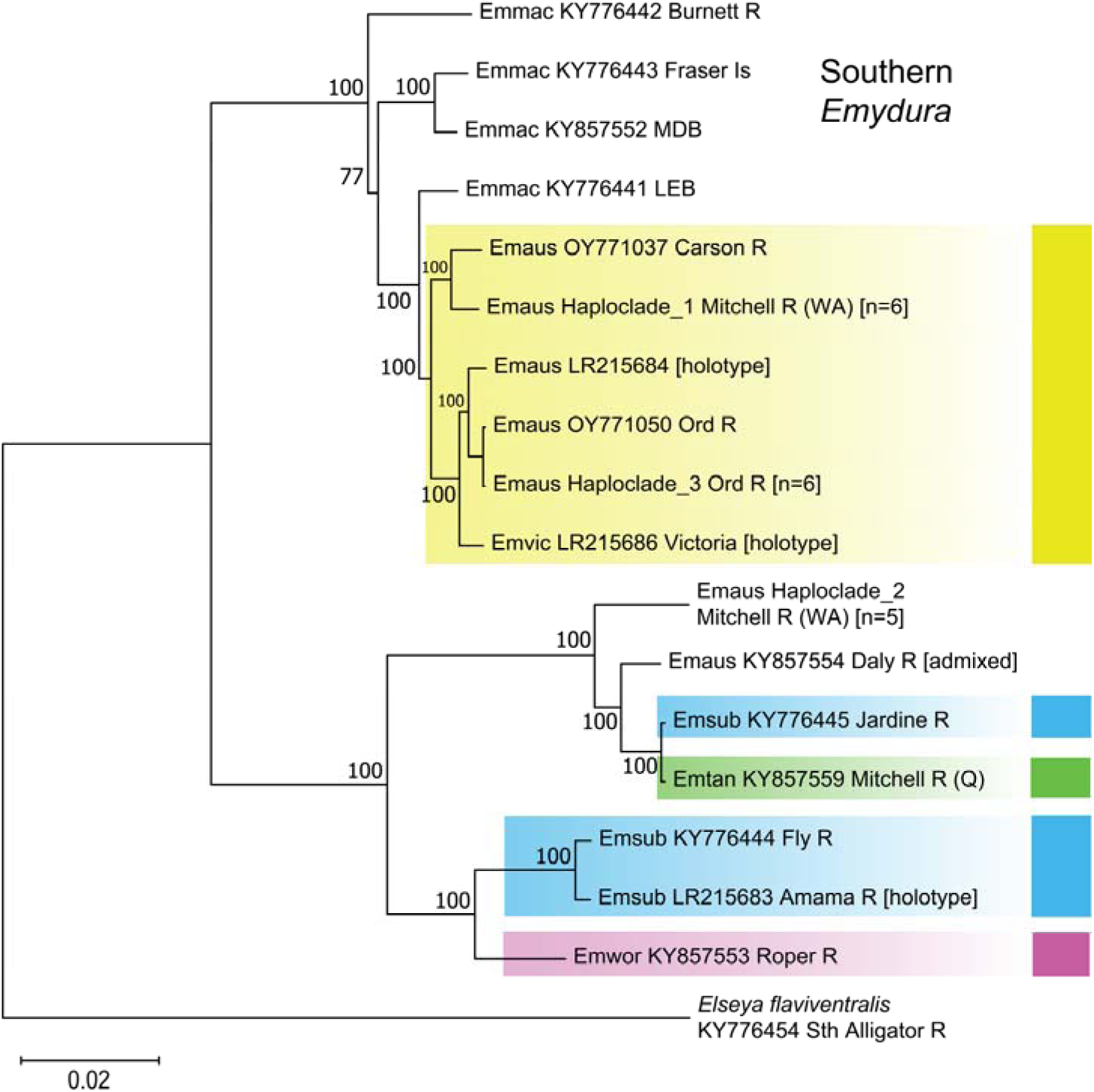
Maximum likelihood phylogeny for the northern *Emydura* based on whole mitochondrion genome sequences. Terminal names comprise the species abbreviation, NCBI accession number and the drainage from which the specimen was sourced (Table S3). Haploclades 1-3 are aggregates of haplotypes that differ by only 5 or less bp (out of 15,984 bp); refer Table S3 for haplotype composition. Coloured bars refer to the diagnosable entities identified in the fixed difference analysis. Note that the Jardine River *Emydura subglobosa subglobosa* individual (KY776445) has the *Emydura tanybaraga* haplotype despite being assigned to *Emydura subglobosa* (blue bars) by the nuclear markers. Tree is rooted with the mtDNA genome of *Elseya flaviventralis* (after Kehlmaier *et al*. 2019). Southern *Emydura* are included for completeness. Bootstraps are based on 1,000 replicates. Refer Table S3 for details of specimens examined.

#### Step 6. Species delineation

At this point of the analysis, we have two independent bodies of evidence, the aggregations of populations into diagnosable entities based on fixed allelic differences and the phylogenies based on sequence divergences. The final step in the analysis is to determine which lineages in the phylogeny, notwithstanding their high bootstrap support, are diagnosable. Here we match the diagnosable groups established in the fixed difference analysis with lineages in the phylogeny to identify diagnosable lineages. Substantial diagnosable aggregations that correspond to lineages in the phylogeny are candidate species. In this case, we have the somewhat unremarkable result that the four substantial diagnosable lineages correspond to the four currently recognised taxa, namely *E. australis*, *E. tanybaraga*, *E. subglobosa subglobosa* and *E. subglobosa worrelli* (Figure 5) The fifth diagnosable lineage, representing the population of *Emydura* in the Mitchell River of the Kimberley escarpment (WA), is enigmatic. Although only four individuals were available to us, the number of fixed allelic differences between this putative taxon and other members of *Emydura australis* was statistically significant, albeit marginally (p = 0.0491, Table 3). The putative taxon included in the nuclear DNA analysis clearly falls within the *E. australis* clade (Figure 5).

This population was not particularly distinctive in the PCA plots (Figures 3 and 4). It was unusual in that 89 loci failed to call for any of the four individuals, whereas that occurred in only 1-10 loci for any of the other populations. The *F_IS_* value for the population was -0.06172, which suggests it is outbred, whereas all other populations had positive *F_IS_* values, ranging from 0.00439 to 0.11074 (mean 0.04224 ± 0.003921SE, Figure S1). The individuals were small and possessed the expanded mouth plates diagnostic for *E. australis* (Georges and Thomson 2010); one of the four had a yellow facial stripe running from the nose to above and behind the eye, whereas the other three had the typical red eyestripes.

## 4 DISCUSSION

### Species delimitation

In this paper, we have brought together the identification of lineages via phylogenetic analysis with analysis at the level of population genetics to yield diagnosable aggregations of sampled populations. In bringing the notion of diagnosability in the form of fixed allelic differences, we were able to identify those phylogenetic lineages on independent evolutionary trajectories. The approach to fixed difference analysis applied here admits the possibility that ends of a geographic cline may be strongly diagnosable, but brought together by sharing of alleles across intervening populations (even a ring species would emerge as a single diagnosable unit). We avoided the possibility of detection of false entities that could arise through patchy sampling (sensu Marshall *et al*. 2021) by comprehensive sampling across the landscape. Although not evident in the results presented in this paper, our approach deals with the uncomfortable fact that reproductive incompatibility is somewhat loosely connected to lineage divergence. A species may emerge as diagnosable from one of many isolates of a species (“budding speciation”) leaving lineages as remaining isolates of the parental species in paraphyly (Avise and Wollenberg 1997, Funk and Omland 2003). These lineages of the parental stock may be subject to sufficient geneflow as not to accumulate diagnostic differences (i.e remain as one “plesiospecies”, Olmstead 1995). Indeed, most mechanisms of speciation will result in paraphyletic taxa as long as reproductive isolation forms the basis for species definition (Patton and Smith 1989). Strict adherence to a lineage concept of species would have those relictual parental clades as cryptic species. Applying diagnosability based on fixed allelic differences to such a scenario would clearly identify that budding speciation had occurred and would inform judgement on the delineation of species within the relictual lineages of the parental species.

We acknowledge that phylogeny is and is likely to remain the cornerstone of future genome-based species discovery but argue that phylogeny should not be the sole or primary focus in species delimitation (Adams *et al*. 2022). Our focus is on species delimitation. Adding the criterion of diagnosability in a suitable framework as we have outlined, informs decisions as to which lineages deserve recognition as valid candidate species versus those that should continue to be regarded as substantive lineages within species, subject to further evidence of speciation becoming available. In addition, speciation does not have to involve the progressive genetic divergence in allelic composition until finally speciation occurs. Positive and negative contributions to the speciation process can arise through belated and often episodic hybridization and introgression between incipient species (Grant and Grant 2019), via the chance assembly of existing genetic variants into novel combinations (Marquis *et al*. 2019) or rapidly via chromosomal rearrangements coming to fixation well before substantial allelic differences arise (Auvinet *et al*. 2018). Neither a phylogenetic analysis based on genome-wide sequence variation nor a search for diagnostic aggregations based on such sequence variation will distinguish species that have emerged recently though discrete processes such as a chromosomal rearrangement leading to reproductive isolation if insufficient time has elapsed to allow appreciable sequence divergence. Some biological species will remain undetected by the approach we apply here.

For the reasons outlined above, we do not want to overstate the value of our approach. It delivers a subset of substantive lineages within a phylogeny (4 or 5 in our case) that can be considered putative species or putative diagnosable lineages within species. The question remains as to whether a substantive lineage is a species or a taxon classified at a lower level (e.g. an Evolutionarily Significant Unit -- ESU) or simply a substantive lineage within species. Our approach is thus not a panacea, but does reduce dramatically the number of otherwise well-supported lineages that would otherwise remain under consideration. With comprehensive geographic sampling, our approach allows more explicit studies of geneflow at the boundaries of putative species, to yield more nuanced, and ultimately more defensible, conclusions on taxonomic status. Indeed, our approach may place a check on taxonomic inflation that has plagued the southern *Emydura* (Georges *et al*. 2018) and other systems (Isaac *et al*. 2004, Frankham *et al*. 2012, Chan *et al*. 2017, Garnett and Christidis 2017, Sukumaran and Knowles 2017, Dissanayake *et al*. 2022).

We embarked on this study anticipating that we would clarify species boundaries within northern *Emydura* and elucidate the historical relationships among them. Past morphological studies, albeit with inadequate quantitation, have identified multiple species and subspecies, including *Emydura australis* (Gray 1841) from the rivers draining the Kimberley Plateau to the north and west and *E. victoriae* (Gray 1842) from the Ord River of Western Australia draining the Kimberley Plateau to the east in Western Australia, the Victoria River and the Daly River in the Northern Territory. The two are variously considered to be one species (*E. australis* has precedence) (Georges and Thomson 2010) or two (Cann and Sadlier 2017). Our results support a single taxon (Figures 5-7) consistent with the findings of Kehlmaier *et al*. (2019), or more strictly, do not provide defensible evidence for two taxa, with the possible exception of the population from the Mitchell River above the Kimberley escarpment discussed in more detail below. *Emydura tanybaraga* was identified in earlier studies based on allozyme electrophoresis (Georges and Adams 1996), with samples from disparate localities in the lower Daly River of the Northern Territory and the Mitchell River of Queensland. Difficulty in distinguishing *E. tanybaraga* from other species because of extraordinary variation in shell shape and facial coloration within species (Cann and Sadlier 2017) resulted in considerable confusion in the distributional range of this species. We have clarified the distribution of *E. tanybaraga* using genetic identification, to show its distribution remains disjunct (Figure 2), and identified boundary drainages (Daly River in the west; Flinders River in the east) where there is contemporary hybridization and introgression. *Emydura subglobosa subglobosa* is a substantive diagnosable lineage found across southern New Guinea and at the tip of Cape York in the Jardine River (Figure 2) and *Emydura subglobosa worrelli* is a substantive diagnosable lineage found in the Daly River and rivers of Arnhem Land above the escarpment, and in rivers draining into the Gulf of Carpentaria east to the Flinders River (Figure 2). This distinction was not evident in an earlier study based on allozyme markers (Georges and Adams 1996), an indication of the greater sensitivity of the SNP analyses. On the basis of the distinction from the other taxa demonstrated in the nuclear marker phylogeny we formally elevate these subspecies to full species status, *Emydura subglobosa* (Krefft 1876) and *Emydura worrelli* (Wells and Wellington 1985), a change from Georges and Thomson (2010) but in line with classifications published elsewhere (Cann and Sadlier 2017).

Hybridization and associated introgression also presents particular challenges for those working with morphological data. The ability to detect hybrid individuals could be of enormous benefit in future morphological studies, permitting the detection and removal of individuals whose morphological attributes are a consequence of their hybrid origin. Undetected hybridization and introgression can confound morphological discrimination between species and render difficult the formal diagnoses and description of species. *E. tanybaraga* from the Daly River is a case in point. How does the morphology of the Daly River *E. tanybaraga* compare to other populations of the species, and how much of the distinction has arisen from the introgression of the *E. australis* genotype in many individuals assigned to *E. tanybaraga*? Indeed, there is a strong possibility that the holotype of this species, from Policeman’s Crossing on the lower Daly River (- 13.767087,130.709915), is admixed with *E. australis*. The ability to assess hybrid and other admixed individuals will provide greater confidence in the reliability of species diagnoses and descriptions in future. We believe that the framework of relationships we present is an excellent platform for assessing and describing the extent of intraspecific morphological variation within taxa well grounded in molecular evidence, and vis-à-vis, how best to describe a species and the extent of interspecific morphology between it and other taxa. There has never been a better time to commence such morphological studies given the current resolution provided by genetic studies, the collections now available to work on around Australia, and the availability of computing software to analyse the morphological data and ask questions of it.

### Mitochondrial Phylogeny

The mitochondrial phylogeny for *cytB* (Figure 6) is consistent with the phylogenies based on nuclear markers (Figure 5) in the support for *Emydura australis*, *Emydura subglobosa* and *Emydura worrelli* as well-defined species. However, *Emydura subglobosa* from the Jardine River at the tip of Cape York Australia has three mitochondrial haplotypes (n=10) that are embedded in the *E. tanybaraga* clade (Figures 6 and 7). We interpret this as having arisen from a recent hybridization event between *E. subglobosa* and *E. tanybaraga* likely consolidated by a selective sweep (Zhan *et al*. 2004, Rato *et al*. 2013, Morales *et al*. 2015) of the *E. tanybaraga* mitochondria through the *E. subglobosa* population. Such lateral exchange of mitochondria is relatively common in Australian chelids (Hodges 2015, Kehlmaier *et al*. 2019). The 8 haplotypes distributed over 46 individuals from the Daly River (Figure 6), many of which were identified in the field as *Emydura australis*, were related to *E. tanybaraga* haplotypes with strong evidence of admixture among these individuals based on the nuclear SNP markers (Table 1). Hence, the *E. tanybaraga* clade in the mitochondrial clade of the *cytB* tree appears to have been complicated as a consequence of recent and contemporary hybridization and introgression between *E. tanybaraga* and the other two species, as we have proposed for *Emydura* in the Jardine River of Queensland, with the additional possibility of incomplete lineage sorting of the mitochondrial haplotypes in the case of the Mitchell River (WA) and Daly River populations.

Included in this complexity were the four individuals from above Mitchell Falls in the Mitchell River of the Kimberley Plateau. They proved to be unusual in many respects. The four individuals had the expanded triturating surfaces on the roof of the mouth characteristic of *E. australis*, three had red eyestripes characteristic of *E. australis* and one a yellow eyestripe characteristic of *E. tanybaraga*. Unlike all other populations examined in this study (*F_IS_* = 0.004393-0.1107), the Mitchell River individuals were outbred (*F_IS_* = -0.06172). A total of 89 SNP loci were not able to be scored for all four individuals (presumably arising from mutations at the restriction enzyme sizes) whereas the comparable data for all other populations ranged from only 1-10. The Mitchell River individuals fell clearly within the *E. australis* clade in both the SNP and SilicoDArT phylogenies (Figure 5), but within the clade comprising *E. tanybaraga* in the mitochondrial *cytB* phylogeny (Figure 6). Together, these data suggest that the genotypes of our four sampled individuals have arisen from introgression between *E. australis* and a second but unsampled taxon in the Mitchell River. The individual depicted in Figure 1j likely represents that unsampled taxon.

In an earlier study of the enigmatic Mitchell River turtles (Kehlmaier *et al*. 2024) using the same whole mitochondrial DNA data, the two distinct haplotype clades of *Emydura* were clearly evident. However, lack of evidence for nuclear genomic differentiation led them to the unambiguous conclusion that only one, albeit morphologically variable, species lives in the Mitchell River. In the mitochondrial evidence, one clade is unambiguously placed by Kehlmaier *et al*. (2024) with *E. australis* to include individuals with both yellow and red faces, and the other clade fell within an ill-defined clade containing both *E. australis* and *E. subglobosa*. We have shown that the Daly River specimen examined by Kehlmaier *et al*. (UC_0245) is likely admixed with *E. tanybaraga* (Table 1) and likely possesses an *E. tanybaraga* haplotype, and that the Jardine River *E. subglobosa* examined by Kehlmaier *et al*. is *E. subglobosa* on the basis of our multiple nuclear markers (Figure 5) but with *E. tanybaraga* mtDNA (Figure 7). Thus, once you take into account our interpretation of the Jardine River haplotypes and the Daly River haplotypes, the haplotypes of these Mitchell River individuals with both yellow and red faces are likely most closely aligned with *E. tanybaraga*, not *E. subglobosa*. Unfortunately, Kehlmaier *et al*. (2024) do not present data on the key diagnostic feature of *E. australis*, the presence of an expanded triturating surface in the upper mouth, absent in both *E. tanybaraga* (very modest expansion) and *E. subglobosa* (Georges and Thomson 2010). Our results show two distinct mtDNA haplotypes (as did Kehlmaier *et al*. 2024) which in our view would be a polymorphism difficult to sustain in the single restricted population of the Mitchell River above the escarpment, given that mtDNA has an effective population size one quarter that of autosomal nuclear genes and a consequentially has a higher extinction rate through drift even in the absence of selection. There is also the evidence of outbreeding in our nuclear gene analysis. We tentatively conclude that there are two species of *Emydura* in the Mitchell River of Western Australia and that they are subject to hybridization and introgression. The four animals we examined were *Emydura australis* based on their nuclear marker phylogeny and the presence of the diagnostic crushing mouth plates. The other species, as depicted in Figure 1j, we provisionally assign as a relictual population of *E. tanybaraga* on the basis of its outward appearance (petite lower jaw) and mitochondrial haplotype (Figure 7). Clearly, further work is required in this remote Mitchell River drainage in the Kimberley region of Western Australia to resolve this issue.

### 4.3 Concluding remarks

Understanding the evolutionary history of diversifying lineages, spatially and temporally, remains a major challenge in evolutionary biology. We undertook comprehensive sampling across the distributional range of our target taxa, the freshwater turtles of the genus *Emydura* in northern Australia and southern New Guinea. This avoided potential identification of false distinction between lineages arising from gaps in sampling as identified by Marshall *et al*. (2021). We used low-cost representational sequencing to bring the fundamental concept of diagnosability in judgements of which substantive lineages are candidate species. We applied our analysis to populations (=sampling sites), that is, independent of the currently accepted taxonomy. With the possible exception of the turtles in the Mitchell River of Western Australia, the diagnosable aggregations of populations corresponded to the four currently recognised taxa in the nuclear DNA phylogenies. We elevated two of these from subspecies to full species. The sister taxa status of *E. subglobosa* and *E. worrelli* (Georges and Adams 1992) is confirmed. The sister relationship between *E. australis* and *E. tanybaraga* reported by Georges and Adams (1992), but not supported unanimously by their analyses, requires further attention. *Emydura australis* may fall outside a clade comprising *E. tanybaraga* and sisters *E. subglobosa/worrelli* (Kehlmaier *et al*. 2024; our *cytB* analysis, Figure 6). The discordance between the multilocus nuclear and mitochondrial phylogenies we attribute to lateral transfer of mitochondria between lineages otherwise on independent evolutionary trajectories, and/or possibly to incomplete lineage sorting of the mitochondrial genome.

Speciation is a (potentially protracted) process, not an event, which complicates analyses that take a phylogenetic lineage approach to species delimitation. Sukumaran and Knowles (2017) called for new methods that accommodate this complexity in deciding which lineages are species as opposed to structure within species. There have been numerous approaches to deal with these issues (see Table 2 of Singhal *et al*. 2018) and we need a more rigorous framework to assess the taxonomic status of significant lineages uncovered by increasingly sophisticated molecular data and analyses (Sukumaran and Knowles 2017, Singhal *et al*. 2018). One of the few downsides of the age of molecular phylogenetics is that our ability to generate simple gene and lineage bifurcating phylogenies from ever-increasingly complex genetic and genomic datasets has tended to reduce our focus on one of the most fundamental concepts in species delimitation, that of diagnosability.

As such, our approach differs from the conventional lineage approach (de Queiroz *et al*. 1998) in that we place our threshold at the time and place where fixed allelic differences have evolved, rather than at the point of initial lineage divergence, which is always difficult to clearly identify given the often-observed gene tree/species tree disparity (Georges *et al*. 2018). We thus differ in placing greater emphasis on diagnosability (fixed differences) rather than on divergence *per se* (which draws also from allele frequency differences).

The power of our approach lies in the ability to distinguish lineages that are represented by diagnosable aggregations of populations or metapopulations that can defensibly become the focus of attention as putative species or substantive diagnosable lineages within species. The final taxonomic decisions in cases of allopatry still require subjective judgements taking into account all available evidence. However, our five-step strategy adds an additional level of objectivity before those subjective judgements are applied and so reduces the risk of taxonomic inflation that can accompany lineage approaches to species delimitation. We believe that our strategy makes a potentially important contribution to any rigorous framework for judgements on species boundaries.

## Supporting information

Supplemental Table 1: Sample locations, SNP data

Supplemental Table 2: Sample locations, cytB

Supplemental Table 3: Sample locations, whole mtDNA

Supplemental Table 4: NewHybrids analysis

Supplemental Table 5: cytB collapsed haplotypes

Supplementary Figure 1: Heterozygosity and FIS estimates

## ACKNOWLEGEMENTS

We would like to thank those who contributed samples to the University of Canberra Wildlife Tissue Collection, and in particular Erika Alacs, John Cann, Sean Doody, Carla Eisemberg, Damien Fordham, Alistair Freeman, Enzo Guarino, Rod Kennett, H. Bradley Shaffer, Scott D. Snyder, Scott Thomson, Jeanne Young and Matt Young. We also thank Jason Carling and Damian Jaccoud for explaining the DArT analysis pipelines and Ian Smales and Ross Sadlier for comments on the manuscript. The study was conducted with funding and/or logistic support from the then Conservation Commission of the Northern Territory, the Bawinanga Aboriginal Corporation, OilSearch Pty Ltd, Exxon Mobil PNG, Balimo Local Level Government, WWF-PNG, the Hermon Slade Foundation and the Piku Biodiversity Network (PBN). Ian Smales and Ross Sadlier provided suggestions that were adopted on the value of this work in giving impetus and direction to future morphological studies and Steve Donnellan provided comments on the interpretation of phylogenetic analyses.

Author Andzrej Kilian declares a potential conflict of interest in that he is the Director of Diversity Arrays Technology Pty Ltd, the company that generated the SNP, SilicoDArT and *cytB* mtDNA data on a cost recovery basis. The remaining authors declare no conflict of interest.

## DATA AVAILABILITY

All mtDNA sequences used in this study are deposited in GenBank, with accession numbers KY776441-5, KY776454, KY857552-4, KY857559 for the whole mtDNA genomes and accession numbers [to be provided on acceptance] for the *cytB* sequences. The sequence alignments are deposited in Dryad, doi: 10.5061/dryad.ht76hdrt8. The SNP and SilicoDArT data and R script that was used in the analysis are also deposited in Dryad under the same doi.

## AUTHOR CONTRIBUTIONS

Conceptualization: AG, PJU; Data curation: AG, XZ, DSBD; Formal analysis: AG, AK, PJU; Funding acquisition: AG; Investigation: AG, XZ, PJU, YA, DSBD; Methodology: AG, PJU, XZ; Project administration: AG; Resources: AG, YA, DSBD; Software: AG, PJU; Supervision: AG; Validation: AG, PJU; Visualization: AG, DSBD; Writing – initial draft: AG; Writing– review and editing: AG, PJU, XZ, DSBD, YA, AK.

XZ generated the whole mitochondrial sequences and lodged them with NBCI; DSBD was responsible for preparing materials for the *cytB* sequencing. AK guided the analyses to generate the SNP data; DSBD and XZ undertook the laboratory work for the SNP analysis. AG led the field work that contributed the bulk of the samples used in this study. AG and PJU led the progress of the manuscript to mature form, to which all authors contributed.

## LIST OF TABLES

**Table S1.** Specimens and their locations of capture used in the SNP and SilicoDArT analyses. PopCode gives the initial population to which they were assigned (="population"). N is the sample size. Drainages follow those defined by Auslig. (2001). Australian Drainage Divisions and River Basins. Canberra, Australia: Commonwealth of Australia. Specimen identity codes are consistent with the UC Wildlife Tissue Collection (UC<Aus>) where additional metadata can be sourced.

**Table S2.** Specimens and their locations of capture used *cytB* sequencing and phylogenetic analyses. PopCode gives the initial population to which they were assigned (="population"). N is the sample size. Drainages follow those defined by Auslig. (2001). Australian Drainage Divisions and River Basins. Canberra, Australia: Commonwealth of Australia. Specimen identity codes are consistent with the UC Wildlife Tissue Collection (UC<Aus>) where additional metadata can be sourced.

**Table S3.** Specimens and their locations of capture used whole mitochondrial sequencing and phylogenetic analyses. PopCode gives the population to which they were assigned (="population"). N is the sample size. Drainages follow those defined by Auslig. (2001). Australian Drainage Divisions and River Basins. Canberra, Australia: Commonwealth of Australia. Specimen identity codes are consistent with the UC Wildlife Tissue Collection (UC<Aus>) where additional metadata can be sourced. Note that specimen KY776454 is misidentified in Genbank as *Elseya dentata* [as of 30-Mar-2025]

**Table S4.** Analysis to provide evidence of contemporary hybridization and admixture among the five currently recognised taxa *Emydura australis, E. subglobosa subglobosa, E. subglobosa worrelli, E. tanybaraga* and *E. macquarii* [NthQLD]. Refer to Table 1 in the main text for explanation. This file contains the supporting analysis. In the spreadsheets labelled subxaus, worxaus etc, outcomes of five replicate runs of NewHybrids are shown and averaged to yield the final reported posterior probabilities.

**Table S5.** Unique *cytB* haplotypes used in the generation of the phylogeny shown in Figure 6. Specimens bearing those haplotypes are listed together with the drainages from which they were sampled. N is the number of individuals with the listed haplotype. Haploclades refer to Figure 6.

## LIST OF FIGURES

**Figure S1.** Observed heterozygosity (a), estimated unbiased heterozygosity (Nei, 1978) (b) and *F_IS_* for populations of *Emydura* across northern Australia, and for the outgroup taxon *Emydura macquarii*. Heterozygosity is based on sequence tags with polymorphic SNPs and so on that basis is subject to ascertainment bias. Nevertheless, the estimates provide an indication of relative heterozygosity across populations. Note that of all populations, only *Emydura australis* from the Mitchell River (WA) shows signs of outbreeding. Vertical bars are standard deviations.

