## Supplementary Figure 1: Heterozygosity and FIS estimates for "Diagnosability to inform species delimitation for the genus Emydura (Testudines: Chelidae) from northern Australia"

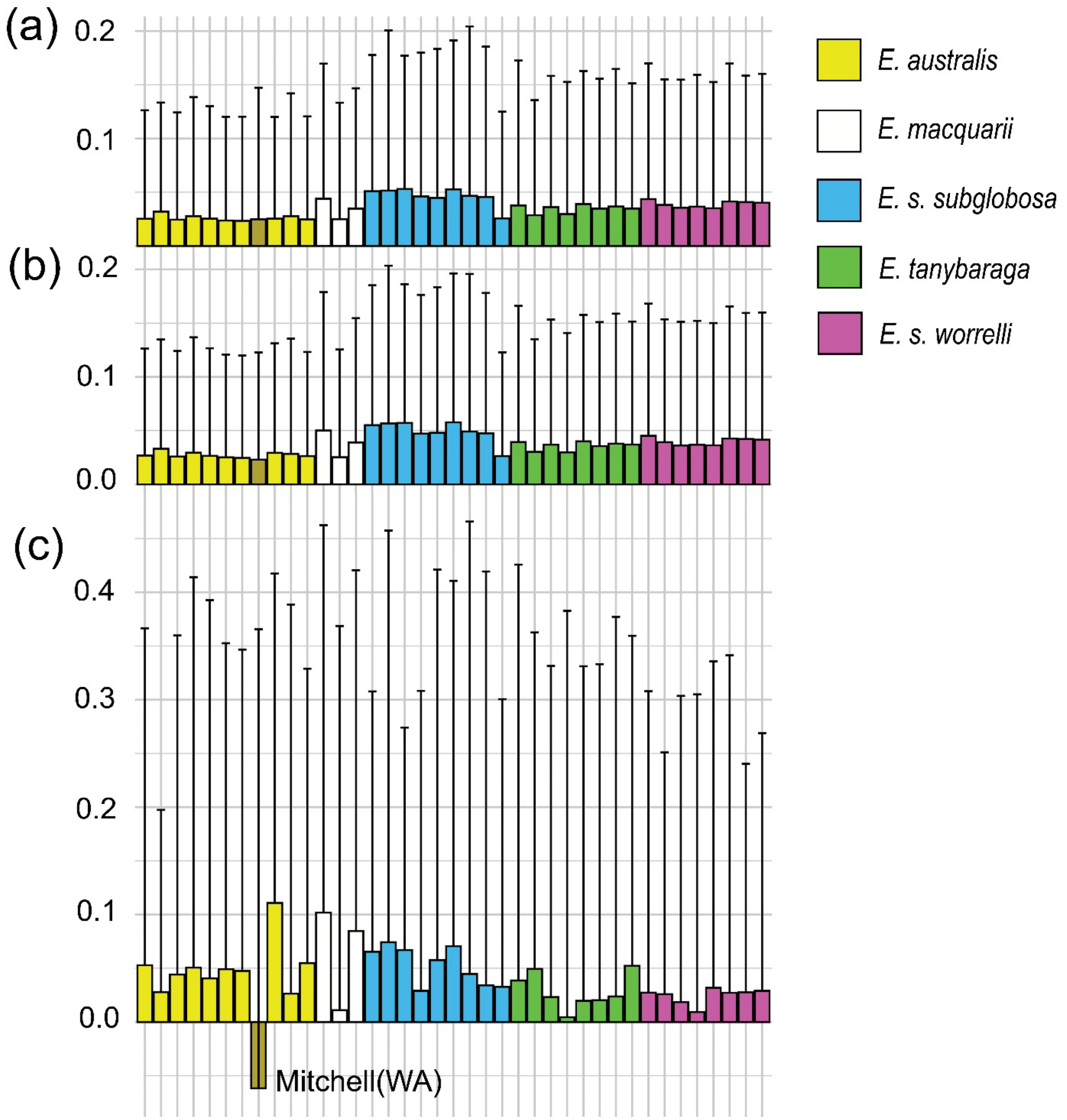


**Figure S1**. Observed heterozygosity (a), estimated unbiased heterozygosity (Nei, 1978) (b) and $F_{IS}$ for populations of *Emydura* across northern Australia, and for the outgroup taxon *Emydura macquarii*. Heterozygosity is based on sequence tags with polymorphic SNPs and so on that basis is subject to ascertainment bias. Nevertheless, the estimates provide an indication of relative heterozygosity across populations. Note that of all populations, only *Emydura australis* from the Mitchell River (WA) shows signs of outbreeding. Vertical bars are standard deviations.
